# Temperature dependence of competitive ability differs from that of growth rate

**DOI:** 10.1101/2022.07.20.499942

**Authors:** Jennifer M. Sunday, Joey Bernhardt, Christopher Harley, Mary I. O’Connor

**Author notes:** Statement of authorship: JS, MO, and CH conceived the experiments, JS and JB collected data, JS designed the experiments, the analyzed data, and wrote the manuscript, and all authors contributed substantially to revisions. Data accessibility statement: Data and code required to reproduce all figures and results will available when the manuscript is accepted.

## Abstract

The effect of climate warming on future community composition is expected to be contingent on competitive outcomes, yet we currently lack mechanistic ecological understanding of how temperature affects competitive ability. Here, we combine resource competition theory with metabolic scaling theory and test hypotheses about how the temperature dependence of competitive ability changes with temperature. We find that the minimum resource requirement for growth, *R*\* – an inverse indicator of competitive ability in phytoplankton – changes with temperature following a U-shaped pattern in all four species tested. The shape of temperature-dependence of competitive ability is systematically different from the temperature-dependence of population growth rates, both in our experiments and in collated data from previous studies. Our results suggest that exploitative competitive success is highest at temperatures that are sub-optimal for growth, and declines rapidly at both cold and warm ends of the thermal performance curve.

## Introduction

Global environmental change is reorganizing biotic communities (Beaugrand *et al*. 2002; Cheung *et al*. 2010). Environmental warming is likely to have greatest effects on biota in the form of indirect effects of temperature, mediated through altered species interactions (Cahill *et al*. 2013). Projecting the indirect, ecological effects of climate change is challenging and requires that we develop our understanding on the mechanistic links between environmental change and rates of ecological processes (O’Connor *et al*. 2015). Such links can produce testable hypotheses for how temperature affects ecological outcomes (Cheung *et al*. 2010; O’Connor *et al*. 2011).

Environmental temperature can affect competitive outcomes, potentially resulting from subtle differences in individual species’ responses in competitive ability with temperature (Birch 1953; Kordas *et al*. 2011; Bestion *et al*. 2018). Changes in competitive outcomes along environmental gradients have been demonstrated in the laboratory and field (Suttle *et al*. 2007; Adler *et al*. 2012; Milazzo *et al*. 2013). Yet despite some theoretical advances, a tested understanding of the mechanistic principles that lead to changes in competitive ability is lacking, limiting our ability to predict ecological outcomes of environmental change (Davis *et al*. 1998; Kordas *et al*. 2011; Alexander *et al*. 2015). Projections of species’ geographic distributions based on large syntheses of thermal performances have advanced our capacity to project climate change impacts (e.g. Pörtner & Knust 2007; Sunday *et al*. 2012), yet these mostly ignore the potential role of competition in limiting species’ occupancy and abundance, particularly at the intermediate spatial scales most relevant to management decisions and the provision of ecosystem services.

Until recently, predictions about how exploitative competitive ability depends on temperature have assumed either that competitive ability is related directly to population growth rate (e.g. Kordas *et al*. 2011; Bestion *et al*. 2018), or that competitive coefficients vary as a direct Gaussian function of temperature (e.g. Urban *et al*, 2012). Although it is straightforward to assume that population growth rate defines competitive ability, which might be appropriate under conditions of high resource supply (Bestion *et al*, 2018), under the resource-ratio hypothesis, competitive ability is attributed to the capacity to grow under very resource-limited conditions (Tilman *et al*, 1982). Thus, the species with a faster growth rate at a given temperature might not necessarily be the most competitive when resources are scarce.

The key to projecting competitive outcomes under competition for a limiting resource is understanding how temperature affects demographic rates at low – i.e. population-limiting – resource levels. When competing species have equal access to a common pool of resources, resource competition theory predicts that the species with the minimum resource requirement will be the competitive winner at population equilibrium (Tilman, 1982). The minimum resource requirement, or *R*\*, is the lowest level of a limiting resource that supports population persistence, also known as the zero-net growth resource level (Tilman, 1982). The species with lower *R*\* can continue to grow when resource availability is *below* the limiting level of the other species.

Minimum resource requirements are expected to be temperature-dependent. Thomas *et al*. (2017) connected resource-limited population growth with temperature across the entire breadth of a species’ thermal performance curve. Their model combines two general tenets of thermal population ecology theory: first, among ectothermic organisms, population growth rates increase exponentially with increasing temperature and then decline (Eppley 1972; Angilletta 2009), leading to a typically unimodal left-skewed thermal reaction norm (Fig. 1a,b,c). Second, resource levels are expected to affect population growth rates in a saturating positive relationship (known as a Monod relationship; or a type-II functional response, although other relationships are possible Fig. 1a,b,d). By allowing birth rates and/or population growth rates to be affected by resource concentration following a saturating relationship, Thomas *et al*. (2017) predicted that an organism’s zero-net growth resource isocline (*R*\*) has a U-shaped relationship with temperature (Fig.1a,b,e), with a shape that differs from the growth rate thermal response curve. Under equal access to a single limiting resource, this relationship provides a prediction of how outcomes of exploitative competition at equilibrium change as a function of temperature (Fig.1f).

**Fig. 1.**
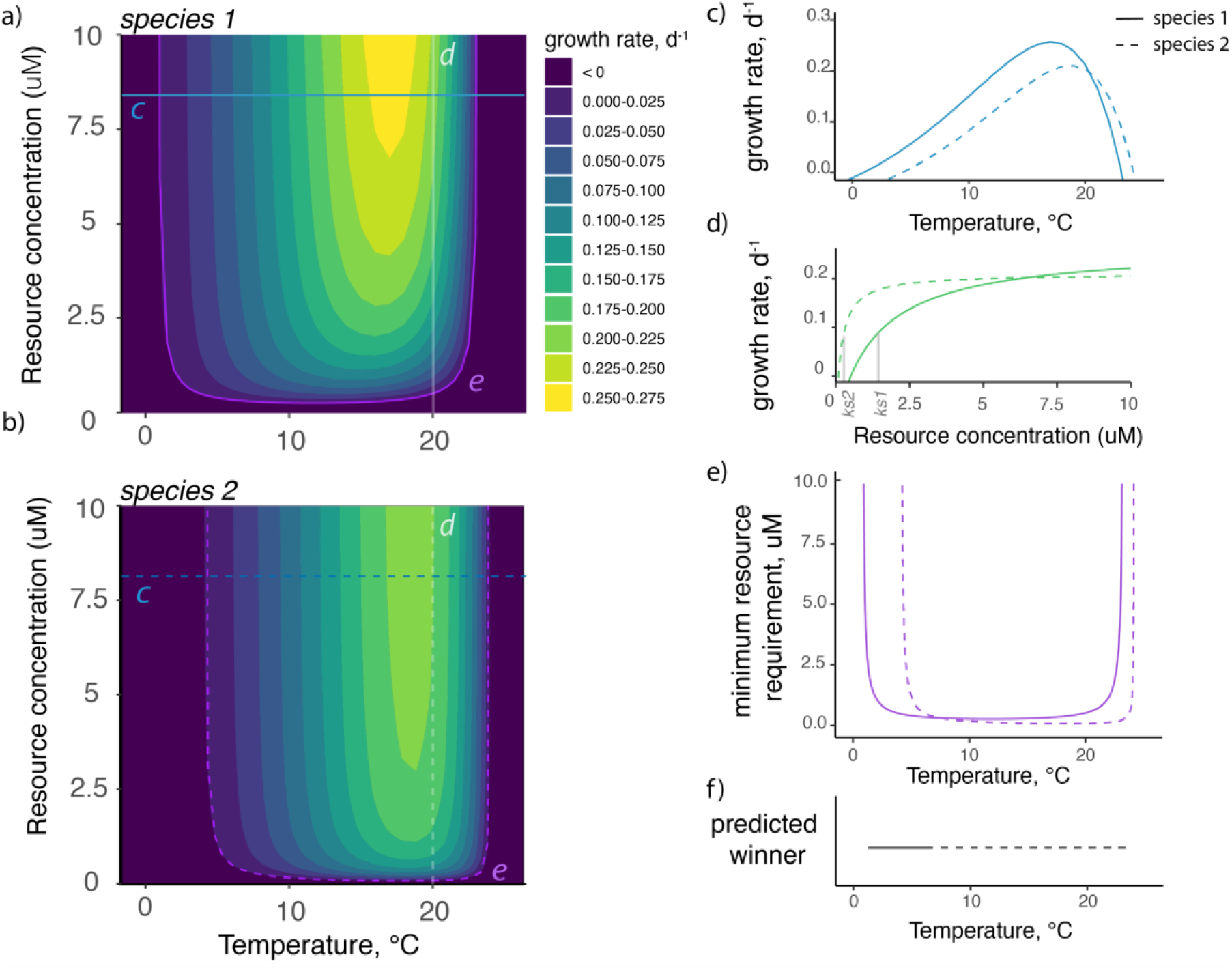
Conceptual relationships between population growth rate, temperature, and resource concentration, with predicted competitive outcomes. Population growth rate relationships with temperature and nutrient concentration are shown for two hypothetical species (a and b). Cutting across a nutrient concentration (blue lines, *c*) gives thermal responses curves for growth (shown in panel c). Cutting across a temperature (white lines, *d*) gives a Monod curve (shown in panel d). Extracting the zero-net-growth isocline as a function of temperature (purple line, *e*) shows the minimum resource requirement for growth, or *R*\* (shown in panel e), which provides a prediction that species 1 will win in a resource competition at low temperatures and species 2 will win at high temperature (f).

Previous empirical work appears to be in line with predictions generated from the Thomas *et al*. (2017) framework. In two species of phytoplankton, Tilman (1982) showed that *R*\* was insensitive to temperature across most of the temperature range with the largest increases at the cold edge of the temperature spectrum in one freshwater phytoplankton species, and at the warm edge in another, together being consistent with a possible U-shaped relationship. *R*\* also increased at cool temperatures in heterotrophic bacteria, sometimes quite dramatically (Reay *et al*. 1999). However, these provide only weak tests of the Thomas *et al*. 2017 model because only parts of the response in *R*\* across temperature were observed, typically below the temperature optimum for population growth. More recently, minimum resource requirements for light and nitrate concentration were found to change with temperature with a pattern that showed, across four temperatures and four species of microalgae, an intermediate thermal optimum in *R*\* (Lewington-Pearce *et al*. 2019). Empirical evidence has also suggested that the thermal optima of species’ thermal response curves decline under nutrient scarcity (Thomas *et al*. 2017; Huey & Kingsolver 2019). While these observations are in line with theory, more precise estimates of curve shape are needed for quantitative tests of theory and comparisons to population growth rates curves.

Here we provide an empirical test of the hypothesis that a species’ thermal performance curve for population growth rate differs from that of competitive ability, assessed as the minimum resource requirement (*R*\*). We take two approaches to empirically quantify the effects of temperature on growth rate and *R*\* in four phytoplankton species, with nitrate as the limiting nutrient. We first build an empirically-grounded prediction of how *R*\* should change with temperature by fitting models of temperature- and resource-dependent population growth to experimental observations across a range of nutrient concentrations and temperatures. From these fitted models we estimate the zero-net growth isocline (*R*\*) as a function of temperature. In our next experiment we allow populations to reach dynamic equilibrium conditions in a semi-continuous flow system, and we measure environmental nutrient concentration at population carrying capacity as a more direct estimate of *R*\*, across temperature treatments (experiment flowchart Fig. S1). We compare all results to test the hypothesis that thermal performance of growth rate is mismatched from thermal performance of *R*\*, by comparing curve shapes and thermal optima. Finally, we draw together previous estimates from the literature of how *R*\* and population growth rates vary with temperature to seek generality in patterns. Our findings provide the most robust empirical support to date to support specific patterns in the temperature response of competitive ability.

## Methods

### Cultures

We used four cultured strains of marine phytoplankton obtained from the Canadian Centre for the Culture of Microorganisms: *Amphidinium carterae* (AC), *Chlamydomonas sp*. (CH), *Chroomonas salina (CS), and Tetraselmis tetrahele* (TT). Each strain was originally isolated in locations off the coast of British Columbia (Table S1), and maintained at 16°C with a 16:8 light:dark cycle under nutrient and light-saturated conditions before experiments.

#### Experiment 1: Estimating population growth as a function of temperature and nutrient concentration

We observed cell density across time during the exponential growth phase under a fully-crossed design of nine nitrate concentrations (0, 11, 22, 33, 55, 110, 220, 330, 550uM), and six temperatures (13, 16, 19, 22, 25, 28°C), in each species. Prior to experimental culture inoculation, we raised subcultures of each algal species for one week in nitrate-free medium to reduce internal cellular nitrate stores by transferring 2ml of high-density culture into 25ml of 0-nitrate medium. See Supplementary Methods for details of nitrate-controlled seawater medium preparation. After this starvation period, each experimental culture was inoculated with 1000 cells ml^−1^ into 12ml medium, kept at constant temperature in one of six incubators (Panasonic MR 154). Light conditions were kept at a 16:8 light:dark cycle. Cell concentration was estimated in each batch culture between 1 and 3 hours after initial inoculation. After this count, cultures at 13°C and 16°C were sampled every 3 days, cultures at 19°C and 22°C were sampled daily, and samples at 25°C and 28°C were sampled twice per day, to estimate changes in cell density during exponential growth. We measured cell densities and biovolumes from 250 uL samples (2% of the initial culture volume) using an imaging flow cytometer (FlowCAM, flow rate = 0.3 ml/min; Fluid Imaging Technologies). This sample volume (2%) was found in previous simulations to improve growth model fits to a greater extent than replicating cultures (Palamara *et al*. 2014). We prepared one culture for each nitrate and temperature treatment level, gaining statistical power through having many levels and fitting nonlinear models (see analysis section). We staggered experiments for each species across time but performed all experiments sequentially within 5 weeks to maximize similarity in culture conditions.

#### Experiment 2: Directly estimating R* across temperature

In a separate experiment, we quantified nutrient concentration at population dynamic equilibrium under semi-continuous flow conditions (see Supplementary Methods) across six temperatures for all four species. We approximated the zero-net-growth resource level, *R*\*, as the asymptotic nitrogen concentration in the medium as the population reached carrying capacity.

Experimental temperatures spanned a wider range of temperatures (3, 10, 17, 24, 31, 38°C) than in Experiment 1 to capture the extreme temperatures of species’ thermal breadths. To create nitrate-limited conditions at carrying capacity we used artificial seawater medium with a reduced nitrate concentration of 4.5 uM (decreased 55-fold relative to standard). Each culture was inoculated with ~250 cells/mL, into 20 mL of nitrate-depleted medium, in a 35ml test tube. We maintained constant temperature and light conditions within six incubators.

We maintained cultures with daily removal of 10% (2ml) of the culture and addition of 2ml fresh sterile nitrate-depleted medium (semi-continuous flow) until equilibrium conditions were met, as assessed visually using plots of cell density and cell size as a function of time. Beginning on the day of inoculation and continuing weekly for the duration of the experiment, cell density and nitrate concentration were measured in each culture (Supplementary Methods). We estimated cell concentration and cell biovolume using a FlowCAM, and assayed nitrate concentration using a cadmium reduction method (LaMotte Nitrate Nitrogen Test Kit) with a Turner Designs Trilogy fluorometer (see Supplementary Methods), which we calibrated with a standard curve before each day of analysis. Although the detection limit of this system is reported as 3uM, our standard curves show that concentrations could be reasonably detected lower than this, to 1uM (Fig. S2). As expected, equilibrium conditions took longer to reach in colder experimental conditions, therefore most replicates lasted 6 weeks minimum and spanned upwards of 10 weeks in 3°C conditions (Figs. S5-S6).

### Statistical analysis

#### Experiment 1. Estimating growth rate as a function of temperature and resource concentration

To test possible formulations of how population growth rate varies with nutrient concentration and temperature, we fit six different models and compared them to identify the best model (Table S2). Among these models were two ‘multiplicative models’ (in which growth rates predicted by a thermal performance curved are reduced multiplicatively to a nutrient-limited growth rate, see Supplementary Methods), and one ‘ interactive model’ (in which nutrients were modeled only to affect birth rates but not death rates, *sensu* Thomas *et al*. 2017, see Supplementary Methods). For the multiplicative models, we used the Norberg model and the Double Exponential model of growth rate as a function of temperature (Table S2), whereas for the interactive model we used only the Double Exponential model (Table S2, Thomas *et al*. 2017). However, we included two versions of each of these three models, one in which the nutrient concentration required to reach half the maximum nutrient-saturated rate of growth (*Ks*, the half-saturation constant of resource-limited growth rate, Fig. 1d, Table S2), was fixed, and one in which *K*s was allowed to increase with temperature. We note when *Ks* is allowed to change with temperature, all models become interactive.

We fit these six growth rate models directly to cell density estimates for the exponential growth phase only. We did this by excluding cell density values at later dates that were *lower* than the exponential-growth trajectory based on earlier dates (Fig. S3-S6). We also identified and removed a few cell density outliers from further analysis (Fig. S3-S6). We used a model fitting approach that gains statistical power by sampling levels along the treatment gradient (e.g., N, T), and estimating confidence through bootstrapping, and then comparing estimates to make inferences.

#### Model fitting

We used the *NLSmultistart* package in *R* (Padfield and Matheson, 2018) to fit each of the six models to our data, whilst exploring a range of starting conditions, and identifying convergence on fitted parameters. We used AIC to compare models that best explained our data for each species. For the best model, we bootstrapped model parameter estimates 100 times to generate 95% confidence intervals around each.

#### Predicting R*

For each species and from each model, we estimated the minimum required resource level (*R*\*) for every half-degree of temperature from −3-50°C, by calculating the nutrient concentration at which growth rate was zero, i.e. root of the growth rate expression using modeled parameters (using the *uniroot* function in *R*). We determined 95% confidence intervals by repeating this calculation for every bootstrapped model iteration. We generated model-predicted thermal optima for growth (*T*opt) as a function of nutrient concentration, in order to compare how this relationship differs among models to affect the *R*\* thermal response curve. When comparing *R*\* values between Experiment 1 and Experiment 2, we subtracted 0.1 from the growth rate expression generated from Experiment 1 before calculating the root, to simulate the daily imposed mortality rate of 10% in Experiment 2.

### Statistical analysis, Experiment 2

#### Estimating R*

We estimated asymptotic nitrate concentration from each replicate as a function of time using equation *RD* in Table S2. Although equation *RD* may not capture the mechanistic relationship between nitrate concentration and time, it allows a robust estimate of the horizontal asymptote, the key parameter of interest.

#### Population growth to equilibrium conditions

We estimated how intrinsic population growth and carrying capacity changed as a function of temperature by fitting a logistic growth model (*LG* in Table S2) to cell densities through time. We used a differential equation solver (*fitOdeModel* function with the ‘PORT’ algorithm in the *simecol* package in R). Because of high sampling error at low cell densities, we set the initial phytoplankton abundance to our mean experimental starting conditions averaged across all cultures. We estimated the functional response in *R*\* and growth rate as a function of temperature by fitting generalized additive models to each parameter for each species.

### Synthesis and Inference

#### Assessing functional responses from both experiments

To test the hypothesis that competitive ability has a different thermal performance curve than population growth rate, we compared the functional responses of *R*\* and growth rate across both experiments. We used the growth rate and *R*\* parameter estimates from the best models from Experiment 1, and the GAM fits to the empirical data from Experiment 2. For each response curve, we assessed curve asymmetry and ‘flatness’ using standard metrics of skewness and kurtosis, as usually applied to frequency distributions (Thomas *et al*. 2016, see Supplementary Methods, negative kurtosis denotes a flatter curve). We did this by modeling each response curve as a positive frequency distribution around a mean (see Supplementary Methods). We compared the optimum temperature for growth rate and *R*\* (*T*opt), defined as the temperature of maximum growth rate and minimum *R*\*, respectively.

#### Compilation of R* estimates from previous studies

To test the generality of the temperature dependence of *R*\* and how it might differ from population growth rate, we extracted previously-published estimates from other species and resource types. We searched Google Scholar using the search terms “R-star”, “resource depletion”, “resource affinity”, and “temperature”, and identified studies in which *R*\* was estimated or could be derived at multiple temperatures. In some studies, estimates of *u_max_* and *K_S_* were estimated empirically and *R*\* was estimated by authors using an arbitrary dilution rate (*D*, ranging from 0.08-0.11 d^−1^ across studies) simulating a continuous-flow experiment following the equation *R*\*= *DK_S_*/(*u_max_-D*) (Tilman *et al*. 1981, Descamp-Julien and Gonzales, 2005). This approach was analogous to ours in Experiment 1, except that rather than explicitly modeling the effect of temperature, the minimum resource requirement was estimated at each temperature. In one study where *u_max_* and *K_S_* were estimated, but not *R*\* (Bestion *et al*. 2018), we used the values reported to estimate *R*\* for *D*=0.1 d^−1^ using the equation above. In other studies, *R*\* was estimated based on resource concentrations remaining in growth medium at population equilibrium in flow conditions (e.g. Reay *et al*. 1999, Lewington-Pearce *et al*. 2019), analogous to our approach in Experiment 2.

## Results

In Experiment 1, we found that the Norberg temperature-resource models were the best models, based on lowest AIC values across all species (Table S3, see Table S2 for model descriptions, see Fig. S7 for displays of model fit). For two species (*Amphidinium carterae* and *Tetraselmis tetrahele*), the Norberg temperature-resource model with a temperature-dependency of *Ks* outperformed the model in which *Ks* was fixed, and *Ks* clearly increased with temperature in these species when Monod models were fit separately within each temperature treatment (Fig. S8). In *Chlamydomonas sp*. and *Chroomonas salina*, *Ks* did not increase across the nitrate concentrations examined, and the models with and without *Ks* increasing with temperature performed similarly (delta AIC < 2; Table S3). The interactive double-exponential model with or without temperature dependence of *Ks* –which was unique in allowing resource concentration to affect birth but not death rates– did not rank highly (Table S3). We present all further predictions using the best model for each species (but see Fig. S9 for *R*\* and *Topt* estimates from all models).

Population growth rates varied with temperature (Fig. 2a) and nitrate concentration (Fig. 2b). For species in which the best models included a term to allow *KS* to change with temperature (AC and TT), model-predicted *Topt* declined at lower nutrient concentrations, as indicated by the shift in the thermal optimum toward lower temperatures (black points in Fig. 2b). For species CH and CS, *Ks* estimates were lower than the lowest nutrient concentration used, as indicated by the sharp saturation of the Monod curves in low nitrate conditions, indicating that our experiment lacked power to detect *Ks* changes with temperatures in these two species.

**Fig. 2.**
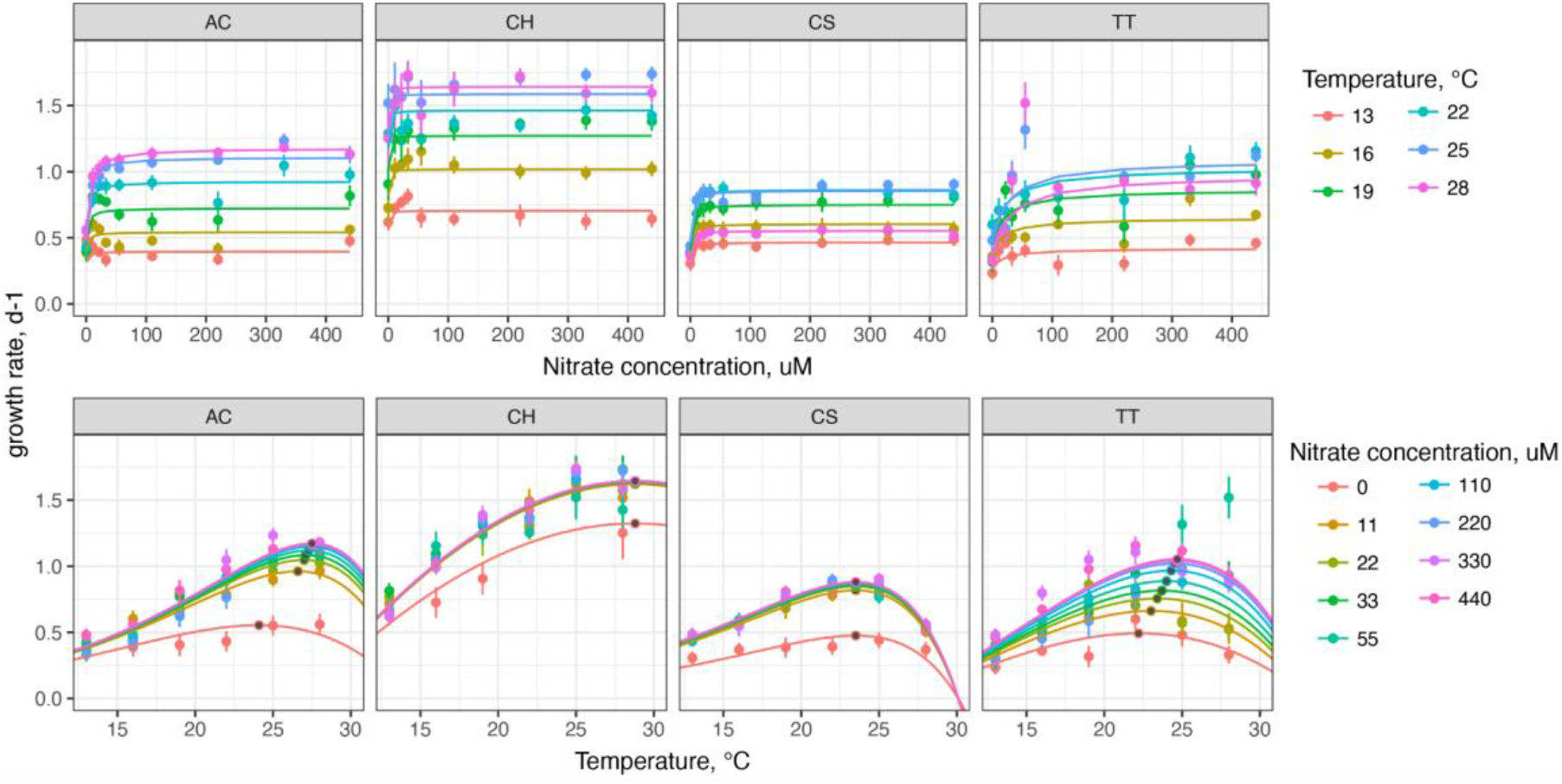
Estimates of population growth rates as a function of temperature and nitrate concentration. Points indicate estimates of growth rates from models fit to cell density data *within* each temperature and nutrient treatment for *Amphidinium carterae* (AC), *Chlamydomonas sp*. (CH), *Chroomonas salina (CS), and Tetraselmis tetrahele* (TT). Lines indicate best-fit model predictions from the best model for each species (Norberg model with *Ks* varying with temperature for AC and TT, Norberg model with *Ks* fixed for CH and CS) that included temperature, nitrate, and time. Black points indicate thermal optimum of population growth rate from best-fit direct models.

The best models (and indeed all of the models, Fig. S9), predicted that minimum resource requirement, *R*\*, would follow a U-shaped relationship with temperature (Fig. 3, lower panels). Notably, *R*\* was predicted to be relatively low across a large range of temperatures, increasing rapidly at cold and warm temperatures where population growth rates approach zero (i.e. the edges of the thermal performance curve).

**Fig. 3.**
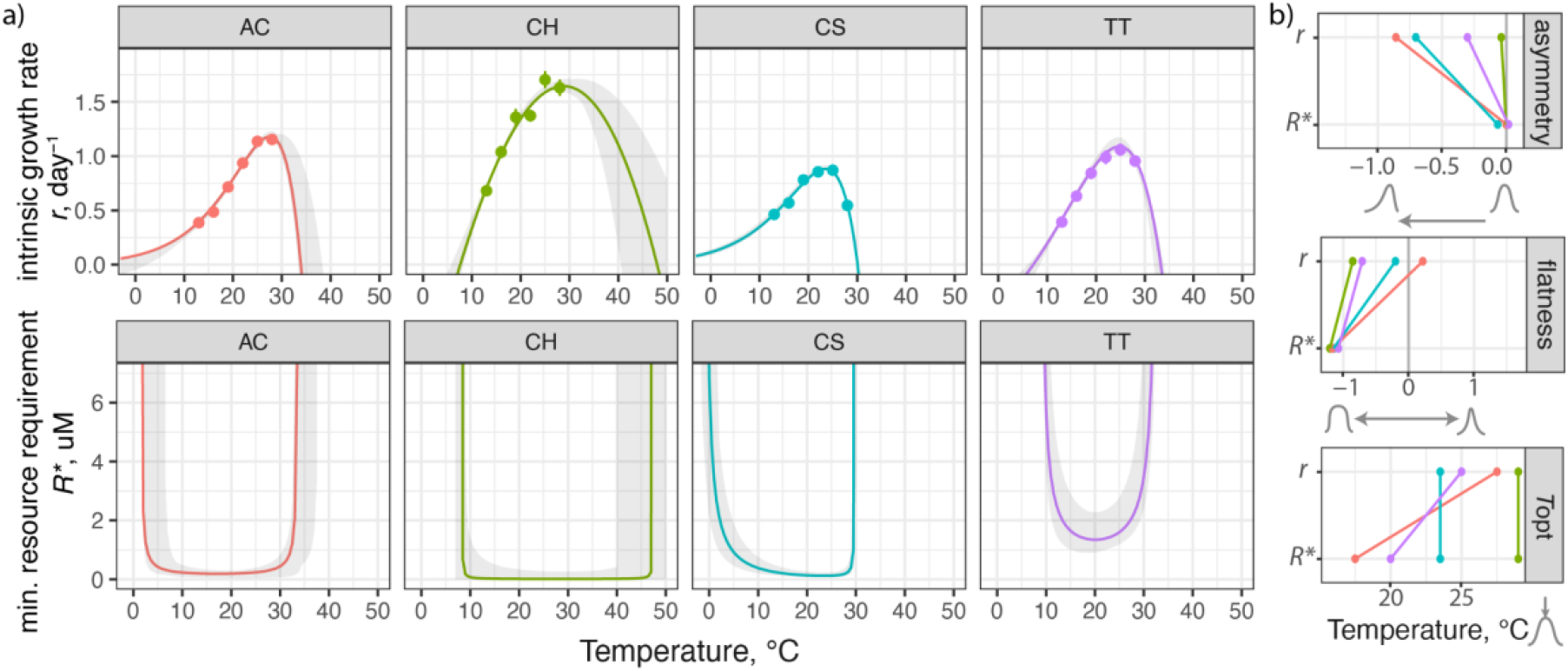
Population growth rates and predicted minimum resource requirement as a function of temperature. (a) Lines represent the mean estimates of nutrient-replete intrinsic growth rate (upper panel) and *R*\* (lower panel) from best-fit model fitted to the data in Experiment 1, for *Amphidinium carterae* (AC), *Chlamydomonas sp*. (CH), *Chroomonas salina (CS), and Tetraselmis tetrahele* (TT). Grey shaded areas represent upper and lower 95% confidence intervals based on model bootstrapping. Points in upper panels indicate estimates of nutrient-replete intrinsic growth rate based on indirect model-fits of growth rate to nutrient concentrations across temperature (fitting a Monod model at every temperature, Table S2). (b) Comparison of curve shape for functional responses of growth rate (*r*) and minimum resource requirements (*R*\*) as a function of temperature of each species. Values represent estimates of curve asymmetry (based on skewness as usually applied to probability distributions), curve ‘flatness’ (based on kurtosis as usually applied to probability distributions) and thermal optima (temperature of maximum *r* and minimum *R*\*). Lines connect values of same species, with color representing species as in panel (a).

In Experiment 2, estimates of *R*\* based on nitrate concentration at population equilibrium were lowest at intermediate temperatures, with a similar an empirically-observed U-shaped relationship with temperature as that predicted by Experiment 1 (Fig. 4, see raw data and model fits in Fig. S10-S11). However, nitrate concentration at carrying capacity was greater than expected at all temperatures when compared to *R*\* predictions of Experiment 1, with minimal concentrations of nitrate nearing 2uM, much higher than the very low values ~0.2uM as predicted from Experiment 1 (see Fig. S12). At the extremely high temperature treatment of 38°C, cells of species AC, CS, and TT died shortly after inoculation (Fig. S10). Although no growth of target cells was observed, final nitrate concentration was lower than that in the exchanged medium (4.5uM), suggesting nitrate might have been consumed by undetected microbes within the experiments. These high-temperature assays were removed from further analysis (shown without removal in Fig. S13). In all cases, a distinct *R*\* minimum was detected in all species, indicating that competitive ability for nitrate reaches a minimum at intermediate temperatures.

**Fig. 4.**
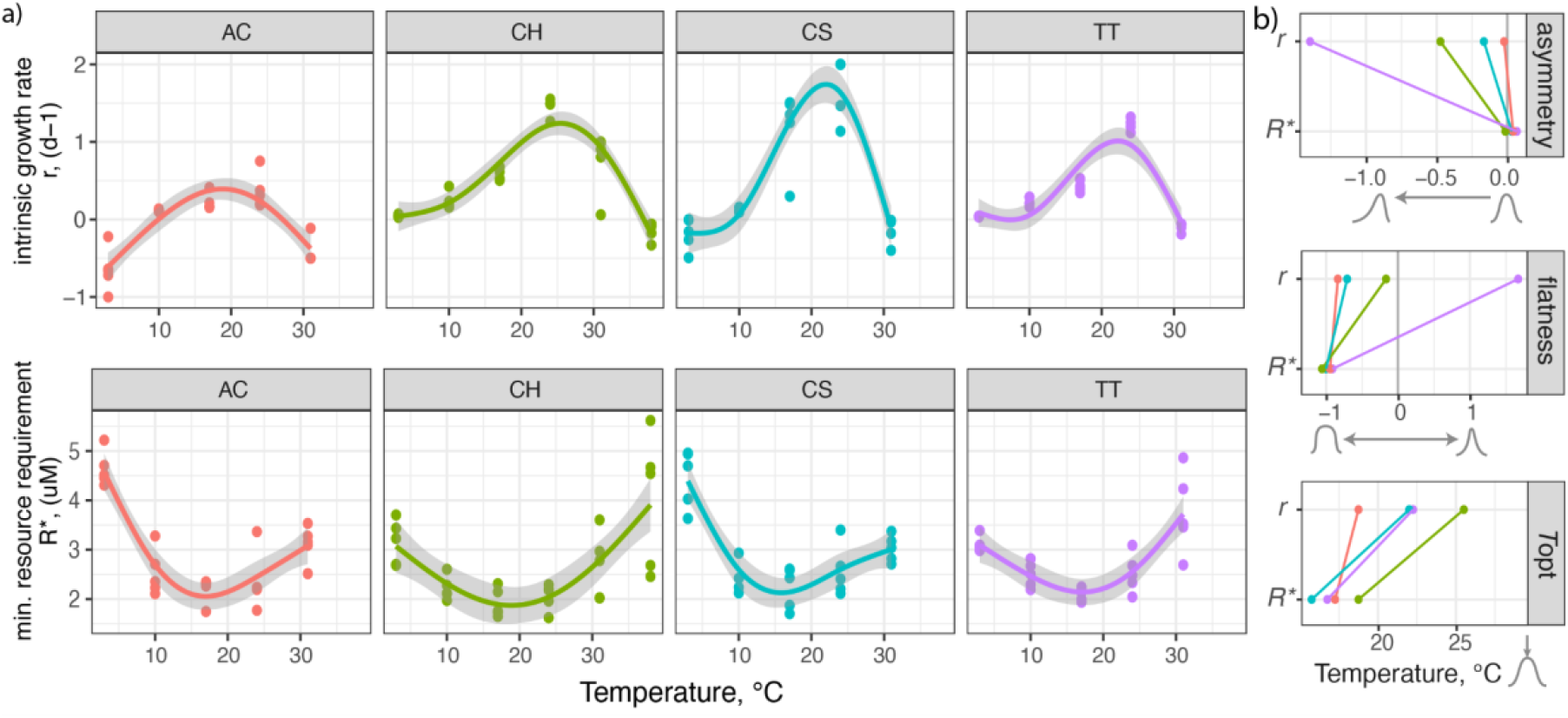
Estimated growth rates (*r*) and minimum resource requirements (*R*\*) from experiments of populations brought to equilibrium under semi-continuous flow. All values from Experiment 2, for *Amphidinium carterae* (AC), *Chlamydomonas sp*. (CH), *Chroomonas salina (CS), and Tetraselmis tetrahele* (TT). Population growth rates (r) were estimated from logistic-growth equations fit to cell density as a function of time; minimum resource requirements were estimated as asymptotic environmental nitrate concentrations at population equilibrium. (a) Points indicate estimated values from each independent replicate, and lines with shading indicate best-fit generalized additive models with 95% confidence intervals. Upper row shows temperature dependance of population growth rate, and bottom row shows temperature dependence of *R*\*. (b) Comparison of curve shape for functional responses of growth rate (*r*) and minimum resource requirements (*R*\*) as a function of temperature of each species. Values represent estimates of curve asymmetry (based on skewness as usually applied to probability distributions), curve flatness (based on kurtosis as usually applied to probability distributions) and thermal optima (temperature of maximum growth rate and minimum R*). Lines connect values of same species, with color representing species as in panel (a).

We found that in both experimental and modeling approaches, the *R*\* curves were more symmetrical (lower skew, i.e. had equal distributions around the mean), flatter (negative values of kurtosis, i.e. changed more slowly around the mean), and had lower optimum temperatures (i.e. minimum *R*\* lower than maximum growth rate) in 6 out of 8 cases, compared to growth curves (Figs. 3b and 4b). This suggests that competitive ability is strong even at cold temperatures for growth, and highest when temperatures are suboptimal for population growth rates.

In our review of previous work, we found five studies in which *R*\* and maximum growth rates had been estimated or could be derived across temperature (Fig. 5). This included 24 unique species and five resource types (silicate, nitrate, ammonium, phosphate, and light) for a total of 37 unique species/resource combinations. With only one exception, estimates of *R*\* were in line with predictions: they either (i) decreased rapidly with temperature at cold temperatures, when population growth rates (*r*) were low, or (ii) were invariant with temperature, typically in the increasing range of growth rate. In the contrary species (*Raphidocelis sp*., from Bestion *et al*. 2018), *R*\* estimates spiked at mid-temperature, completely out of line with predictions (dashed purple line in Fig. 5). With or without including this species, optimal temperatures for maximum growth rates within species/resource combinations were greater than optimal temperatures for minimum *R*\* (paired t-test with *Raphidocelis sp*. included, df = 37, p = 0.00052, difference = 4.1 ± 2.2*_95%CI°_*C; excluded, df=36, p= 0.00011, difference = 4.1 ± 2.1*_95% CI_*°C). Transforming each curve horizontally so that values are shown as a function of the optimum temperature allows across-species patterns to be observed along a common temperature-deviation-from optimum axis. Across species, *R*\* curves transformed this way appear more symmetrical and flatter (leptokurtic) compared to growth rate curves (Fig. 5b).

**Fig. 5.**
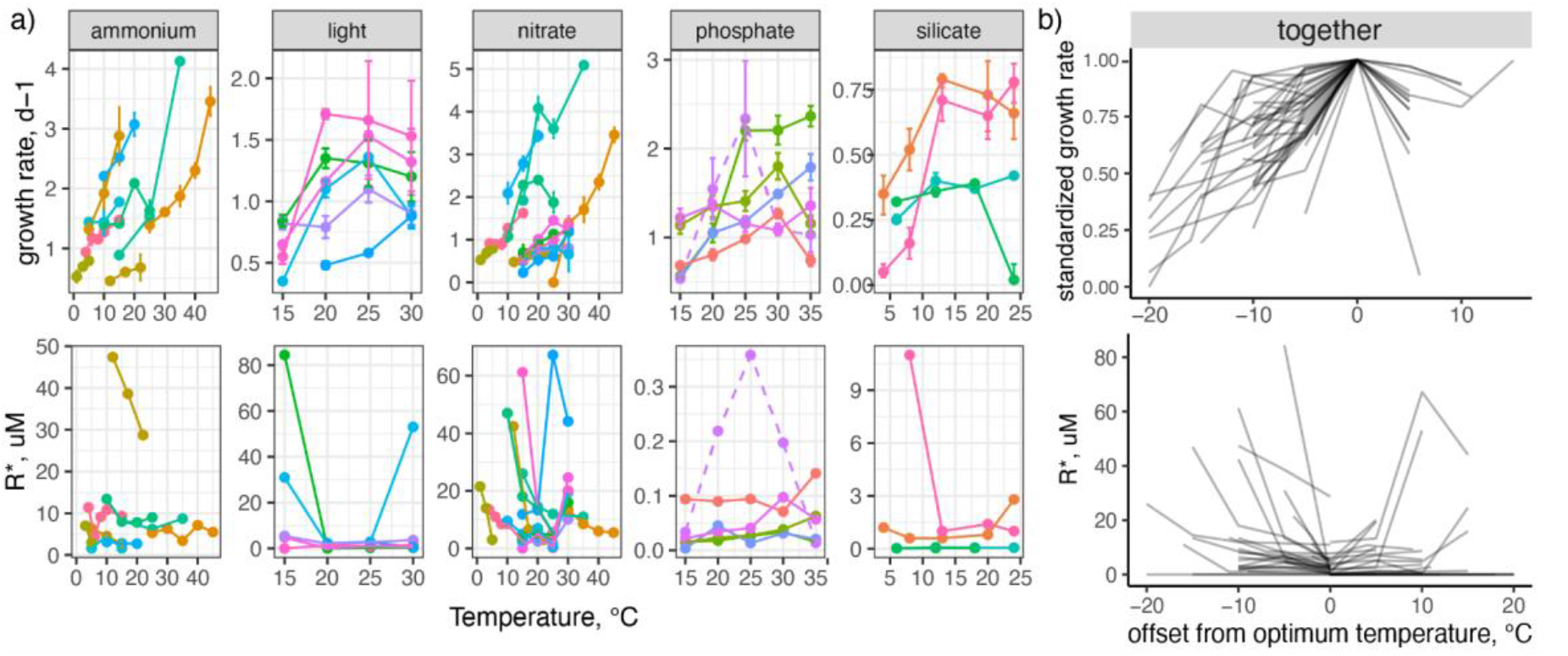
Review of population growth and minimum resource requirement (*R*\*) across temperatures compiled from other studies. (a) Colors represent species of bacteria and microalgae from 5 previous data sources in which growth rate and *R*\* were estimated or could be derived across multiple experimental temperatures (Tilman *et al*. 1981; Reay *et al*. 1999; Descamps-Julien & Gonzalez 2005; Lewington-Pearce *et al*. 2019). Dashed lines highlight a single result counter to general predictions. (b) Same results shown after standardizing temperatures to degrees (°C) above or below the optimum temperature for each metric; lines have slight transparency to show overlap. Growth rates were standardized across species to indicate growth rates relative to within-study maximum (hence all curves cross 0,1).

## Discussion

These results provide empirical evidence that the minimum resource requirement (*R*\*) changes with temperature in a manner that differs from classical left-skewed growth rate thermal performance curves. *R*\* typically has a lower temperature optimum, a wider breadth of high performance, and a more symmetrical decline toward temperature extremes, compared to thermal performance curves for population growth rate in the same species. We assessed these relationships in two ways and four species, providing independent experimental evidence across species and methods. Our work corroborates previous theoretical predictions (Thomas *et al*. 2017) and empirical findings of a mid-temperature minimum in *R*\* (Lewington-Pearce *et al*. 2019). The finding that competitive thermal performance curves (i.e. the inverse of *R*\*) differ from growth-based thermal performance curves has strong implications for how we expect outcomes of resource competition to change across thermal regimes. Here we discuss the combined effects of nutrients and temperature on growth rate, and the implications of these findings for understanding how competitive dominance is altered by temperature.

Our results lend support to previous findings that the optimal temperature for growth declines with nutrient concentration (the metabolic meltdown hypothesis, Thomas *et al*. 2017; Huey & Kingsolver 2019) in two ways. First, for two species in Experiment 1 the best-fitting model was that in which *Topt* changed with temperature (black points in Fig. 2, Fig. S9). Second, in our empirical observation of *R*\* (Experiment 2), the optimum temperature for *R*\* was lower in all species compared to the *Topt* for growth at higher resource concentrations (Fig. 4 and 5). Hence, as nutrients were lowered, the temperature at which there is *any* growth (defacto *Topt* at lowest resource concentrations), was lower than the temperature at which there is optimal growth at high resource concentration. These results support the metabolic meltdown hypothesis: when resources are limited, thermal optima decline (Huey & Kingsolver 2019). More broadly, a decline in *Topt* with resource concentration has also been observed in two microalgae (Thomas *et al*. 2017), macroalgae (Gerard 1997), salmon juveniles (Brett *et al*. 1969; Brett 1971), and cod (Peck *et al*. 2003).

Our results also suggest that the half-saturation resource constant, *Ks*, is temperature-dependent in at least two species, based on model fits in Experiment 1. In the other two species, our nitrate treatments were probably too high to accurately detect changes in *Ks* since these species reached maximum growth rates even in lowest non-zero nitrate concentration of 11um (Fig. 2). An increase in *Ks* with temperature matches theoretical predictions (Goldman & Carpenter 1974; Aksnes & Egge 1991; Reuman *et al*. 2014). We note this does not mean that resource affinity itself declines with temperature, indeed we would expect *K*s to increase in proportion with maximum resource-saturated growth rate (*u*max) should specific resource affinity remain constant with temperature (Reay *et al*., 1999). Given the mixed evidence for a relationship between *K*s and temperature in previous work (reviewed in Bestion *et al*. 1999) and the importance of this relationship for the shape of competitive changes with temperature, our work underscores the need for a better understanding of mechanistic temperature-dependence of resource assimilation mechanisms.

Although changes in minimum resource requirements (*R*\*) with temperature were similar in pattern across our two experiments, the predictions in Experiment 1 were lower than the values in Experiment 2 in all but one species (*Tetrahele tetraselmis*). This offset could be attributed to a few mechanisms. First, in Experiment 1 we might have underestimated *R*\* because we lacked power to parameterize the decline of growth rates when nutrient concentration was low. Since growth was fairly high to saturating at our lowest non-zero experimental nutrient concentration (11uM) in all species but *Tetrahele tetraselmis*, we could have overestimated *Ks* and therefore *R*\*. Second, nitrate concentration at equilibrium might have been overestimated in Experiment 2 if cells had burst during filtration, leading to increased nutrient concentration in the assay medium. Third, there could have been evolutionary change between or during experiments (conducted one year apart), although this is unlikely because maintenance culture conditions matched those of their source culturing facility, and adaptation during the longer second experiment would be expected to reduce, not increase, the minimum nitrate requirement for growth. Fourth, final nitrate concentrations in Experiment 2 might have been lower than our assay system could detect, although our standard curves suggest that nutrient concentrations between 2uM were detectable (Fig. S2). While more precise estimates of *R*\* will be useful for predicting outcomes of competition among specific pairs of species, the changes in *R*\* across temperature within species were consistent across experiments.

Our analysis of other microorganisms suggests that these patterns generalize in microbial aquatic species. The theoretical predictions of a u-shaped relationship between the minimum resource concentration (*R*\*) and temperature matches empirical observations across taxa and resource types. One exception was the single species of microalgae in Bestion *et al*. (2018) which had an intermediate maximum of *R*\*. Perhaps this is a species with a different physiological response to temperature, or perhaps this was due experimental error: within this same study, the fit of the Monod curve to the growth rate data within temperatures was especially poor for this species (Bestion *et al*. 2018). Nevertheless, our results indicate that thermal optima for competitive ability are generally lower compared to growth rate when data exist

We interpret the minimum resource requirement for growth, *R*\*, as being inversely predictive of competitive ability, but this requires further empirical testing. Certainly relative *R*\* has been predictive of the outcome of resource competition among organisms for which resources are mixed and equally available among competitors (Tilman *et al*. 1982). The prediction has mainly been supported in micro-organisms in a mixed aquatic environment that compete for dissolved nutrients or light (reviewed in Miller *et al*., 2005). The framework might generalize to motile organisms that compete for moving prey; Gilbert *et al*. 2014 conceptualized the minimum resource level for growth in more general predator prey systems as *R*\*=m/ea (where m=mortality rate, e=conversion efficiency, and a=attack rate). Applying metabolic scaling theory, mortality (Gillooly *et al*. 2007) and attack rates (based on changes in metabolically-mediated velocities of consumers and resources; (Dell *et al*. 2014)) can each be predicted to change with temperature (Gilbert *et al*. 2014). Changes in the relative velocities of consumers and resources with temperature therefore likely play a key role in determining the shape of the *R*\* curve with temperature in unmixed predator-prey systems, and deserves further investigation. Indeed, temperature-dependent swimming speed appears to be key to competitive outcomes in Dolly Varden and white-spotted char in stream experiments (Watz *et al*. 2019; Yamada *et al*. 2020). We note that our results do not likely generalize to non-motile species competing for locally drawn-down resources (i.e. plants).

We present evidence that thermal performance curves for minimum resource requirements (*R*\*) differ in shape from thermal performance curves for growth rate, with greater competitive abilities at cooler temperatures than might be expected based on growth rate alone. This puts into question the idea of predicting species distributions based on intrinsic rates of growth across temperature (i.e. the fundamental thermal niche), and provides a mechanistic approach for predicting geographic ranges based on temperature and competitive environments, when resource competition is thought to be important (i.e. the realized niche). Using thermal performance curves of minimum resource requirement (*R*\*) as an indicator of competitive ability and therefore occupancy might be most relevant for systems in which resources have sufficient time to be drawn down to levels that exclude species. The temperature-dependance of *R*\* might similarly be useful for predicting how resource-driven succession will be altered by temperature (e.g. Tilman, 1990). By contrast, relative population growth rate might be a more useful predictor of competitive success where there is high turnover and instability. Finally, the different shapes of *R*\* thermal performance curves compared to growth rate thermal performance curves might reconcile why thermal performance curves of growth often don’t predict species’ distributions at geographic scales (Araújo *et al*. 2013). Even if species have similar upper thermal tolerance limits (Sunday *et al*. 2011, 2019; Araújo *et al*. 2013), competitive ability at temperatures cooler than the thermal optimum for growth might dictate their success outside of the tropics (Fig. 6). Overall, refining our understanding of the interconnections between temperature, competitive ability, and population growth in realistic multi-species assemblages will advance a mechanistic understanding of species distributions, biodiversity changes across temperature gradients, and response to climate change.

**Fig. 6.**
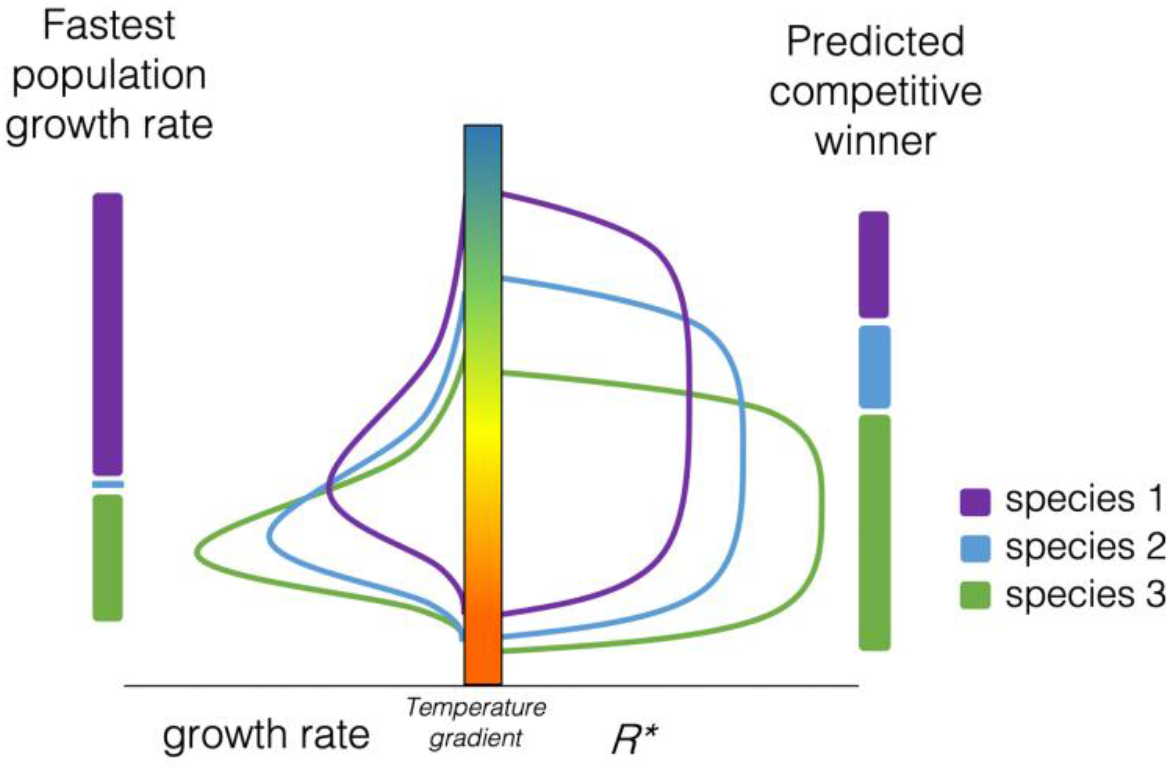
Predicted distributions based on expectations of competitive ability along a temperature gradient are different from predicted distributions based on population growth rates. Left hand side shows resource-replete growth rates for three hypothetical species, with predicted winners based on fastest growth rate. Right hand side shows the minimum resource requirement for growth and the predicted competitive dominant in equilibrium conditions.

## Acknowledgements

We are grateful to a team of dedicated research assistants for culture maintenance and experimental assays: Scott Brydle, William Ou, Kimmy Hofer, Zander Chilla, Michelle Nguyen, Amir Gohari, Jane Yangel, and Blaire Cameron. We were supported financially by the Natural Sciences and Engineering Board of Canada, a Canada Foundation for Innovation Leaders Opportunity fund grant to C.D.G.H., and specific funding from the Biodiversity Research Centre at the University of British Columbia to JMS. C.D.G.H.

## Notes

### Competing Interest Statement

The authors have declared no competing interest.

