## Supporting materials for "Temperature dependence of competitive ability differs from that of growth rate"

List of supporting materials:

Supplementary Methods
Supplementary Citations

Table S1. Names and collection locations of four marine phytoplankton species used throughout study.
Table S2. Summary of equations and models used.
Table S3. AIC scores of the six models fit to all 4 species.

Fig. S1. Flow chart of Experiments 1 and 2 used to test predictions about R* and growth rate changes with temperature
Fig. S2. Standard curves for nitrate analysis
Fig. S3. Cell counts of *Amphidinium carterae (AC)* as a function of time
Fig. S4. Cell counts of *Chroomonas salina "var. 1" (CS)* as a function of time.
Fig. S5. Cell counts of *Chlamydomonas sp. (CH)* as a function of time.
Fig. S6. Cell counts of *Tetraselmis tetrahele (TT)* as a function of time.
Fig. S7. Displays of fit for models of cell density as a function of time, temperature, and resource concentration in Experiment 1.
Fig. S8. Patterns of $K_S$ with temperature, from indirect model fitting within each temperature treatment.
Fig. S9. Predicted growth rate, optimum temperature, and minimum resource requirement from each model.
Fig. S10. Logistic growth models fit to cell density as a function of time in Experiment 2.
Fig. S11. Decay function fit to nitrate concentration as a function of time in Experiment 2.
Fig. S12. Minimum resource requirements ($R*$) from Experiments 1 and 2 displayed together.
Fig. S13. Estimated growth rates ($r$) and minimum resource requirements ($R*$) from experiments of populations brought to equilibrium under semi-continuous flow.

### Supplementary Methods

*Details of sub-hypotheses among model formulations*

The modeling framework put forward by Thomas *et al*. (2017) includes a few assumptions that bear empirical testing. First, the manner in which resource concentration affects growth rate – either affecting birth rates only, or affecting both birth and death rates together, affected the shape of the $R*$ response curve with temperature (Thomas *et al.* 2017). Hence the way resource limitation manifests as a change in population growth rate can determine the shape of temperature dependence of competitive ability and the extent to which it differs from temperature dependence of population growth rate. Second, the half-saturation constant in a saturating resource-growth relationship (*Ks*), which describes how quickly growth rates increase with resource concentration, was assumed to not vary with temperature. However, theoretical studies predict *Ks* to increase with temperature (Goldman & Carpenter 1974; Aksnes & Egge 1991; Reuman *et al.* 2014) - meaning that higher levels of resources are needed to achieve half-maximum growth rate in warmer temperatures. Empirical studies find a range of responses on the temperature sensitivity of *Ks* (reviewed in Bestion *et al.* 2018), suggesting there is more to uncover.

*Different combinations of temperature- and nutrient-dependent growth curves*

We began with two previously-proposed models of how population growth rate varies as a function of temperature, which each fit a unimodal growth relationship with variable levels of skew: the *Norberg* model (Table S2) and the *Double-Exponential* model (Table S2, Thomas *et al*. 2017). We included the *Double-Exponential* model because it articulates birth and death rate terms separately, allowing a derivation in which only birth rates are affected by resource concentration (Thomas *et al*. 2017). Following Thomas *et al*. 2017, we assumed that these models estimate population growth rate at saturating nutrient conditions, such that reduced nutrients would reduce population growth rate following a Monod equation (ME in Table 1, Fig. 1). We substituted the Norberg or Double-Exponential models for growth into $u_{max}$ (asymptotic population growth rate at saturating resource concentrations) of the Monod model, producing the Norberg Resource model and the Double Exponential Resource model (Table S2). These represent *multiplicative models, sensu* Thomas *et al.* (2017). We included a third model that allowed only birth rates to vary as a function of resource concentration, leaving death rates independent of resource concentration, reproducing the Interactive Double Exponential model of Thomas *et al.* (2017; Table S2), introduced to avoid the illogical effect of increasing growth rates when resources are low (Thomas *et al.* 2017).

*Allowing half-saturation constant to vary with temperature*

We tested the sub-hypothesis that the half-saturation constant varies with temperature, by exploring versions of each model in which *Ks* was either held constant or allowed to increase with temperature according to the *Ks(T)* equation in Table S1. We chose an exponentially-increasing relationship for *Ks(T)*, which also fixed modeled estimates of *Ks* as positive across temperatures, but note the other shapes are possible.

Previous nutrient-temperature-growth rate models (e.g. Thomas *et al.* 2017) have not allowed for a temperature-dependency of *Ks*, which means (counterintuitively) that specific nutrient affinity changes with temperature (i.e. the initial *slope* by which growth rate increases with nutrient concentration in nutrient-limited conditions, Reay *et al.,* 1999), with the same temperature dependency as maximum growth rate. Our approach thus allowed more variation in how nitrate affinity changed with temperature.

*Model fitting*

We thus produced six models of growth rate as a function of temperature and resource concentration: the *Norberg Resource* model and *Double Exponential Resource* model, and the *Interactive Double Exponential Resource* model, and the counterpart of each in which *Ks* was allowed to vary with temperature. Based on observations of low but positive population growth in our 0-Nitrate treatments, we included a term in all six models, $z$, to represent an undetected source of Nitrate, potentially due to intra-cellular stores, and substituted $R+z$ into all expressions of resource concentration ($R$). This improved all model fits but did not change overall results.

In order to fit each model of population growth rate to our empirical observations of cell density across time, we substituted each expression of growth rate into the exponential growth equation (Table S2), so that models could be directly fit to cell density observations. We thus simultaneously fit cell density to time, resource concentration, and temperature. This *direct* approach has been shown to improve accuracy of parameter estimates compared to fitting a series of models to parameter estimates of previous models (Palamara *et al.*, 2014). Results were similar when models were fit indirectly, and we use indirect models to visualize growth rate estimates within treatments (points representing growth rate in Figs. 2-3).

*Artificial Seawater Medium*

To control nitrate in sterile seawater medium, we prepared *Enriched Seawater, Artificial* Water medium throughout our experiments (ESAW; Harrison *et al.* 1980). For Experiment 1, we first made a single batch of medium that excluded $NaNO_3$, and subsequently adding a dilute $NaNO_3$ stock solution to nine separate batches, following sterile ESAW preparation protocols (Harrison *et al.,* 1980). In Experiment 2, to reduce the chances of small variation in medium conditions among experiments, we prepared one large batch of nitrate-depleted ESAW medium to be used for the entire experiment, which was autoclaved and stored in separate 1L batches.

*Semi-continuous flow conditions*

In Experiment 2, we maintained a semi-continuous flow with daily removal of 10% (2ml) of the culture and addition of 2ml fresh sterile nitrate-depleted medium; hence there was a 10% per day exchange of medium. Cultures were gently aspirated 5 times in order to mix the cultures before culture removal and replenishment. All sterile work was performed in a laminar flow hood using sterile techniques. Once per week a small amount (0.5-1ml) of sterile deionized water was added to the cultures at 38°C and 32°C in order to offset evaporative water loss in these cultures.

*Nitrate assays*

Once per week nitrate concentration was collected from 1.33ml out of the 2ml daily removal of culture medium. This aliquot was syringe-filtered with a GF/F filter, stored at -80°C, and later thawed for nitrate assay. We assayed nitrate concentration using a cadmium reduction method (LaMotte Nitrate Nitrogen Test Kit) and measured absorbance using a Turner Designs Trilogy fluorometer (Nitrate/Nitrite Module, P/N: 7200-074). Assays were conducted on 6 days, and each day we first created a standard curve using serial dilutions of nitrate in fresh nitrate-free medium (Fig. S2).

*Nitrate reading outliers*

In several nitrate readings (n=18, 2% of all readings), the reading was extremely high, exceeding the original concentration of the medium and that of the daily exchange medium, and more than double the concentration in adjacent readings of the same culture tube. We observed no corresponding change in population size, and therefore removed these readings, deeming them to be an artifact of the nitrate estimation method rather than an unaccounted source of nitrate in the system. Results are similar when these outliers were not excluded, but error estimates substantially larger.

*Curve shape parameters*

We assessed the shapes of temperature response curves using parameters that are typically used to describe the moments of central tendency in a frequency distribution. We did this by transforming the model-fit response curves of each model and species, truncated at the lowest and highest observed temperatures of each experiment, into a frequency distribution, following methods of Thomas *et al.* 2016. Because $R*$ is an inverse of competitive ability, and we were interested in comparing competitive ability with growth rate curves, we inverted $R*$ into a unimodal positive curve by subtracting values from an arbitrary nitrate concentration of 6uM. This created a positive unimodal curve comparable to a normal distribution around an optimum. We then calculated kurtosis and skew using standard metrics of density distributions (using the *skewness()* and *kurtosis()* functions in *R*).

We thus used skewness of the frequency distribution as an assessment of curve asymmetry, in which negative values denote a negatively-skewed curve relative to a normal distribution at 0, and positive values denote a positively-skewed curve (see inset images in Fig. 3b and 4b). Similarly, we used kurtosis of the frequency distribution as an assessment of curve 'flatness', in which negative values denote a flatter curve compared to a normal distribution at 0, i.e. values descend more slowly away from the mean, and positive values denote a pointier curve (see inset images in Fig. 3b and 4b).

Table S1. Names and collection locations of four marine phytoplankton species used throughout study.

| **Species name** | **Species code** | **Class** | **Collection location in BC** |
|---|---|---|---|
| *Amphidinium carterae* | AC | Dinophyceae | Bamfield |
| *Chlamydomonas sp.* | CH | Chlorophyceae | Strait of Georgia |
| *Chroomonas salina "var. 1"* | CS | Cryptophyceae | Bamfield Inlet |
| *Tetraselmis tetrahele* | TT | Prasinophyceae | Vancouver Island |

Table S2. Summary of equations and models used. (a) Expressions used to build larger models in Experiment 1 or to estimate equilibrium conditions in Experiment 2, (b) temperature-resource-growth models used in Experiment 1. Multiplicative models are those in which resource level affects intrinsic growth rate in a multiplicative sense, while interactive refers to that in which resource level affects only birth rates, not death rates (Thomas *et al.* 2018).

a)

| **Name** | **Expression** | **Definitions** |
|---|---|---|
| Exponential growth equation | $ln(N(t)) = N_0 + r * t$ | *N*=cell density (ml-1), *r* = population growth rate ($d^{-1}$), *t*=time (d) |
| Monod equation | $r\ (R) \ = u_{max} * \frac{R}{Ks\ +\ R}$ | r = population growth rate ($d^{-1}$), umax = population growth rate at replete resource concentrations, R= resource concentration, Kks=half-saturation resource constant |
| Norberg TPC model | $r\ (T) = a * e^{bT}\left(1 - \left(\frac{T - z}{\frac{w}{2}}\right)^2\right)$ | r = population growth rate at replete resource concentrations ($d^{-1}$), T=temperature (°C), a,b,z,w = model parameters |

| | | |
|---|---|---|
| Double exponential TPC model | $r\ (T) = b_1\ e^{b_2 T}\ -\ (d_0\ +\ d_1\ e^{d_2 T})$ | r = population growth rate at replete resource concentrations, ($d^{-1}$), T=temperature (°C), $b_1$,$b_2$ = birth rate parameters, $d_0$,$d_1$,$d_2$=death rate parameters |
| *Ks* variation with temperature | $K_s(T) = {K_{s0}}^{T+f}$ | *T*=temperature (°C), $K_{s0}$, *f* = model constants |
| Resource decline with time | $R(t) = R^* \left(\frac{t + R_0}{b + t}\right)$ | $R_0$ = starting nitrate concentration (uM), $R^*$ is asymptotic nitrate concentration, $b$ = decay constant, *t* = time (d) |
| Logistic growth | $\frac{dN}{dt} = rN\left(1 - \frac{N}{K}\right)$ | *N*=cell density, *r* = population growth rate ($d^{-1}$), *K* = carrying capacity, *t* = time (d) |

b)

| **Name** | **Expression** | **Definitions** |
|---|---|---|
| Norberg Resource model | $r\ (T, R) = a * e^{bT}\left(1 - \left(\frac{T - z}{\frac{w}{2}}\right)^2\right) * \frac{R}{Ks\ +\ R}$ | Substituting Norberg growth equation for umax in Monod equation |
| Norberg Resource model, *Ks* varies | $r\ (T, R) = a * e^{bT}\left(1 - \left(\frac{T - z}{\frac{w}{2}}\right)^2\right) * \frac{R}{{K_{s0}}^{T+f}\ +\ R}$ | Substituting $K_S(T)$ for *Ks* in above. |
| Double Exponential Resource model | $r\ (T, R) = (b_1\ e^{b_2 T}\ -\ (d_0\ +\ d_1\ e^{d_2 T})) * \frac{R}{Ks + R}$ | Substituting Double Exponential growth |

| | | |
|---|---|---|
| | | equation for umax in Monod equation |
| Double Exponential Resource model, *Ks* varies | $r\ (T, R) = (b_1\, e^{b_2 T}\ -\ (d_0\ +\ d_1\, e^{d_2 T})) * \frac{R}{{K_{s0}}^{T+f} + R}$ | Substituting *$K_S(T)$* for *Ks* in above. |
| Interactive Double Exponential Resource model | $r\ (T, R) = b_1\, e^{b_2 T} * \frac{R}{Ks + R} -\ (d_0\ +\ d_1\, e^{d_2 T})$ | Substituting birth rate component of Double Exponential growth equation for umax in Monod equation, and subtracting death rate |
| Interactive Double Exponential Resource model, *Ks* varies | $r\ (T, R) = b_1\, e^{b_2 T} * \frac{R}{{K_{s0}}^{T+f} + R} -\ (d_0\ +\ d_1\, e^{d_2 T})$ | Substituting *$K_S(T)$* for *Ks* in above. |

Table S3. AIC scores of the six models fit to all 4 species. Emboldened values indicate model within 2 units of the lowest AIC score. See Table S2 and for model descriptions.

| species name | acrn. | temperature-resource model (number of terms) | | | | | |
|---|---|---|---|---|---|---|---|
| | | multiplicative | | | | interactive | |
| | | *Norberg* Ks fixed *(7 terms)* | *Norberg* Ks=f(T) *(8 terms)* | *Double-Exponential* Ks fixed *(8 terms)* | *Double-Exponential* Ks=f(T) *(9 terms)* | *Interactive Double-Exponential* Ks fixed *(8 terms)* | *Interactive Double-Exponential* Ks=f(T) *(9 terms)* |
| *Amphidinium carterae* | AC | 982 | **963** | 984 | 965 | 994 | 975 |
| *Chlamydomonas sp.* | CH | **865** | **867** | 869 | 872 | 870 | 871 |
| *Chroomonas salina* | CS | **706** | **707** | **707** | 709 | 730 | 735 |
| *Tetraselmis tetrahele* | TT | 980 | **976** | 1025 | 1146 | 977 | 987 |

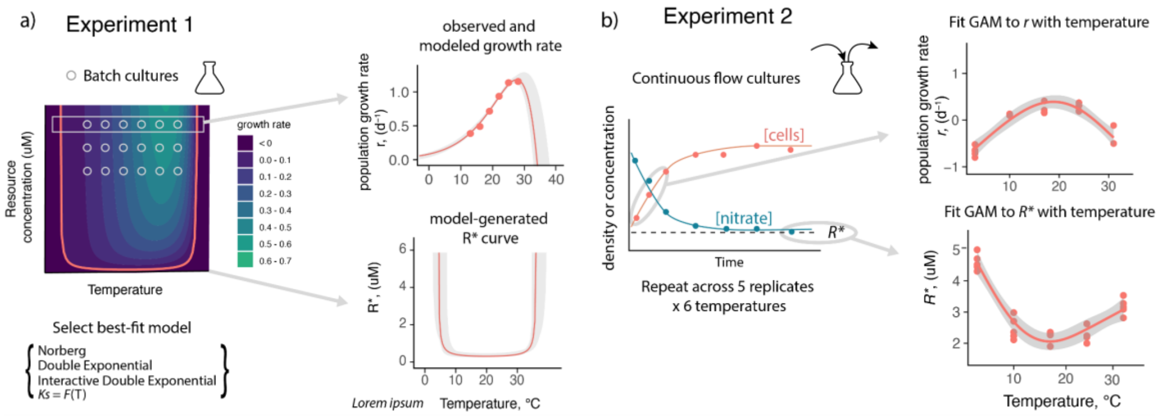


Fig. S1. Flow chart of Experiments 1 and 2 used to test predictions about R* and growth rate changes with temperature. In Experiment 1 (a) we used batch cultures to assess intrinsic rates of population growth at different temperature and nutrient conditions (represented as crosses). We fit six models of population growth rate as a function of temperature and nutrient concentration to these data, and identified the best-fitting model. We extracted predictions of growth rate (r) with temperature at a saturated nutrient condition (top inset in a), and extrapolated $R*$ with temperature by calculating the root, i.e. where growth rate in our models would be zero (lower inset in a). In Experiment 2 (b) we used continuous flow of cultures, and fit models to observations of population growth rate (logistic growth) and of nitrate concentration (decay model) as a function of time. We repeated across 5 replicates and 6 temperature conditions, extracted model-fit parameters of r from logistic growth model (top inset in b) and of asymptotic nutrient concentration, or R* (bottom inset in b), and fit generalized linear models to generate predictions of each as a function of temperature.

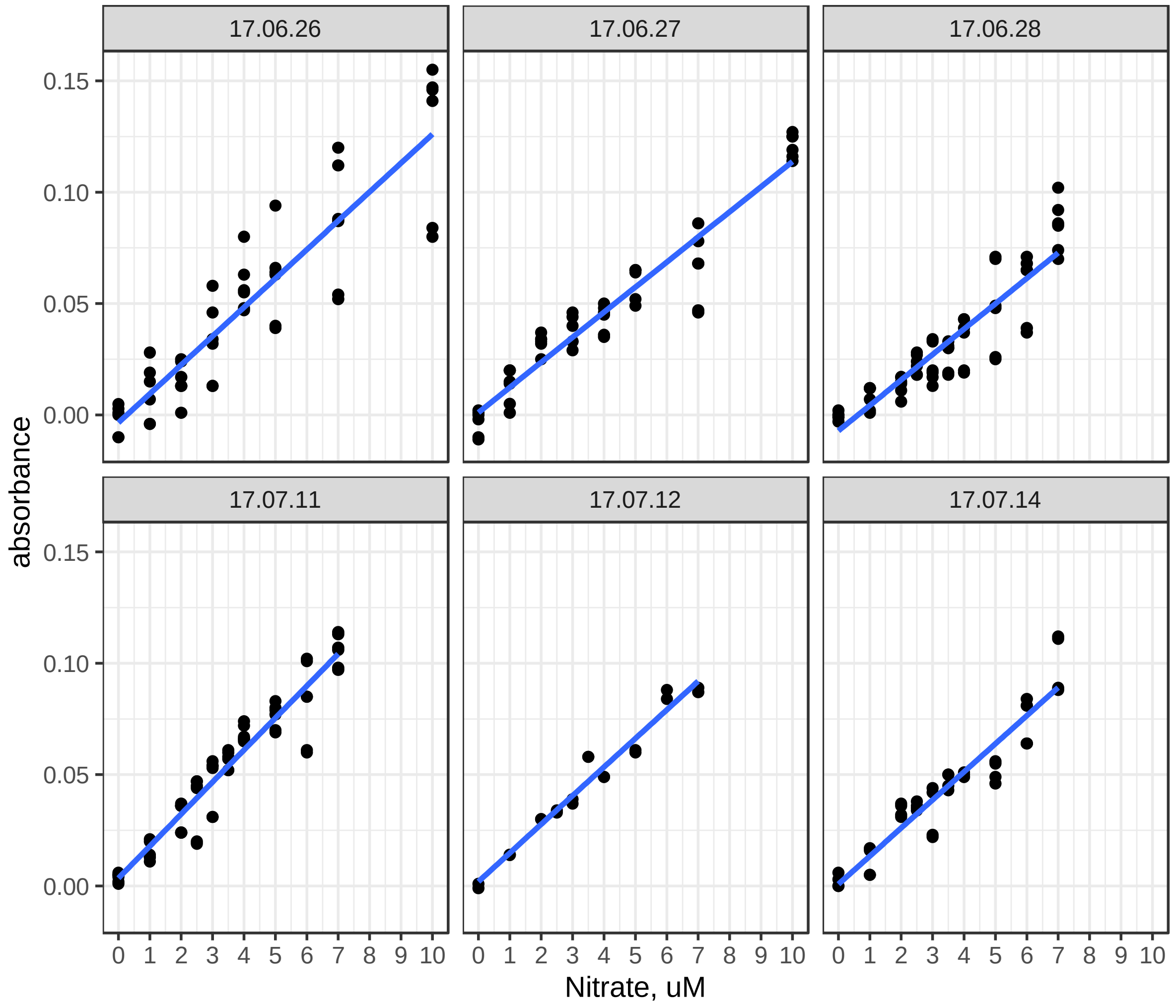


Fig. S2. Daily standard curves for calibration of Nitrate analysis system. Labels denote date (dd.mm.yy), nitrate values are those created using serial dilutions of nitrate-containing medium into nitrate-free medium. Blue lines indicate linear regression of absorbance as a function of nitrate concentration.

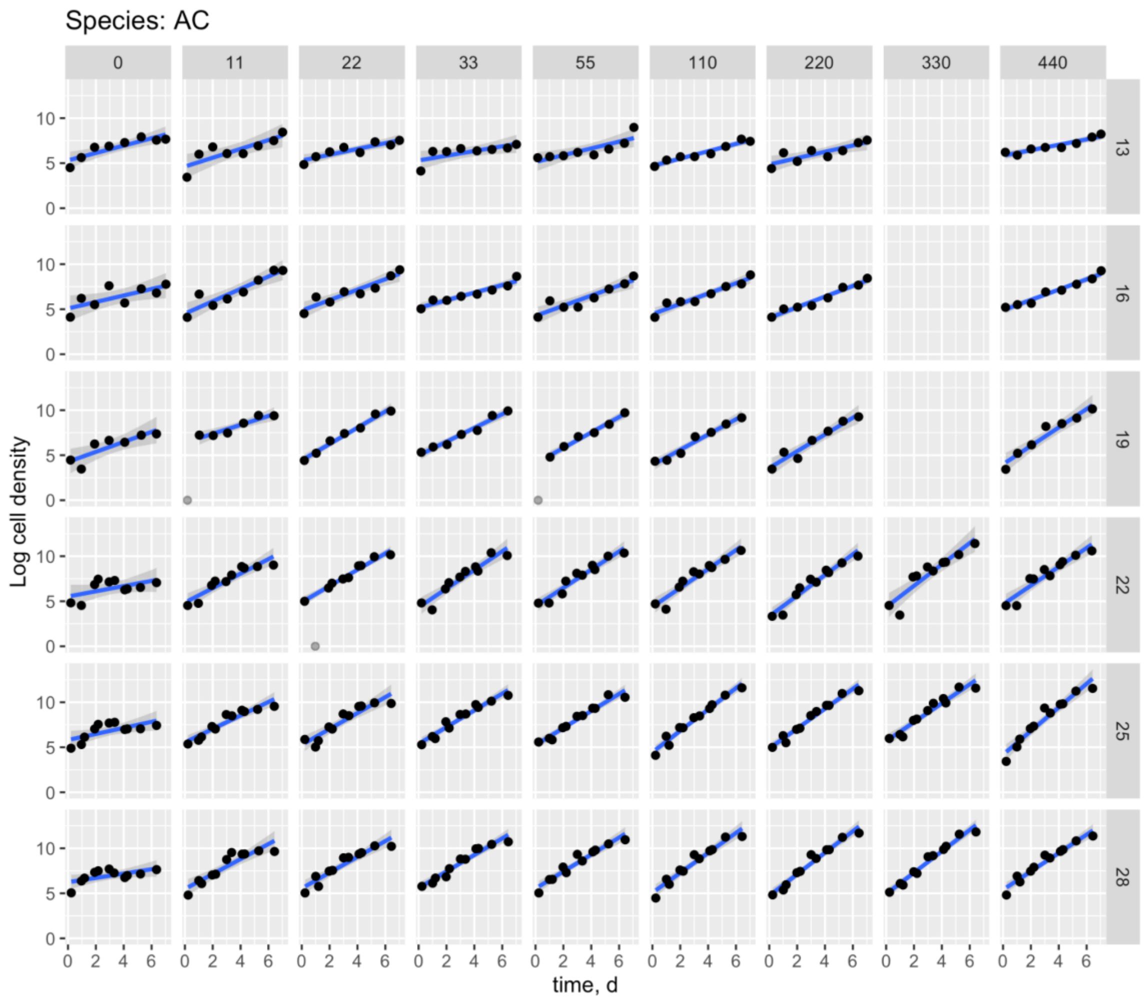


Fig. S3. Cell counts of *Amphidinium carterae (AC)* as a function of time. Grey data points indicate observations putatively outside the exponential growth phase, or deemed as outliers, and left out of linear model fits. Blue lines with shading show best-fit linear model with standard error to logged cell density, representing intrinsic rate of growth (*r*). Three cultures were lost (at 330uM) during the experiment.

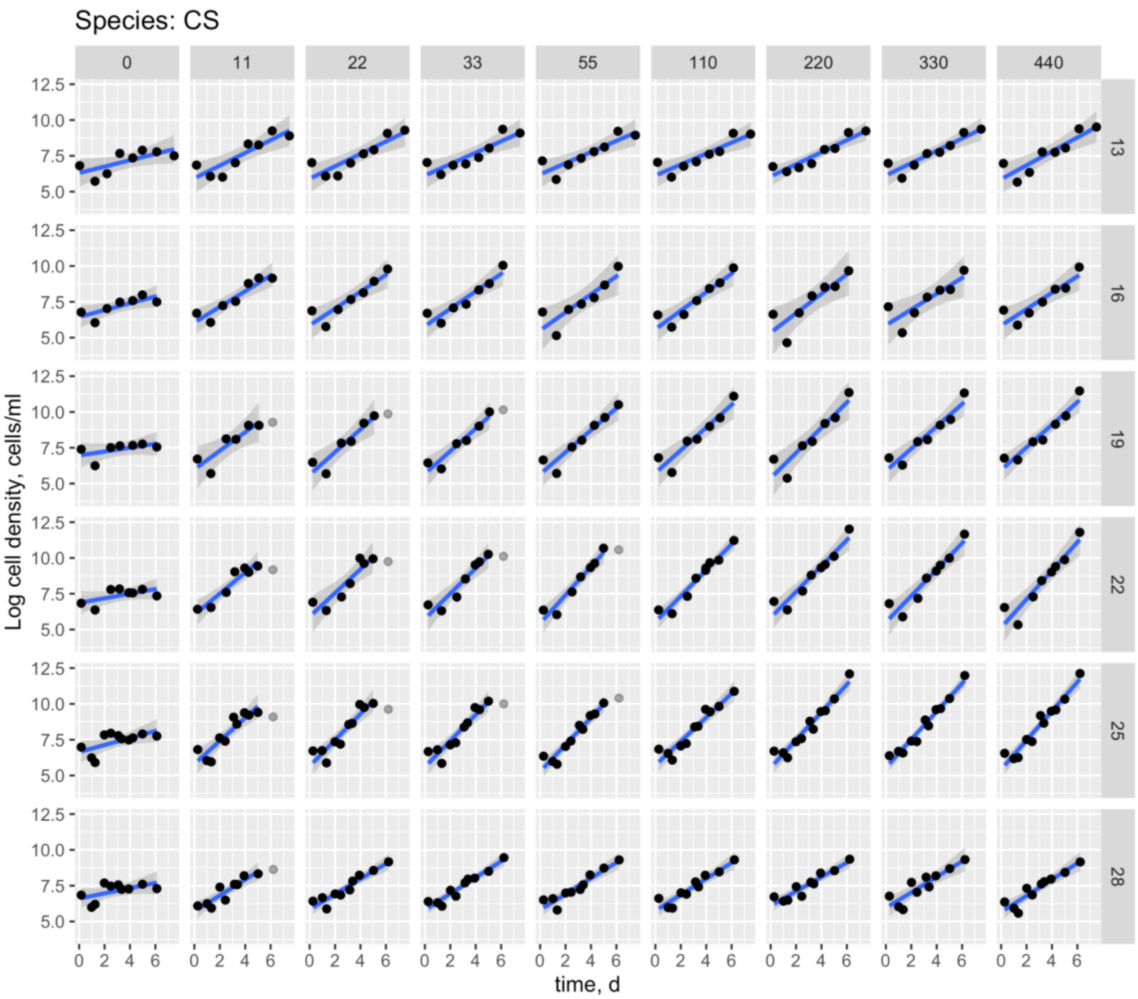


Fig. S4. Cell counts of *Chroomonas salina "var. 1" (CS)* as a function of time. Grey data points indicate observations putatively outside the exponential growth phase, or deemed as outliers, and left out of linear model fits. Blue lines with shading show best-fit linear model with standard error to logged cell density, representing intrinsic rate of growth (*r*).

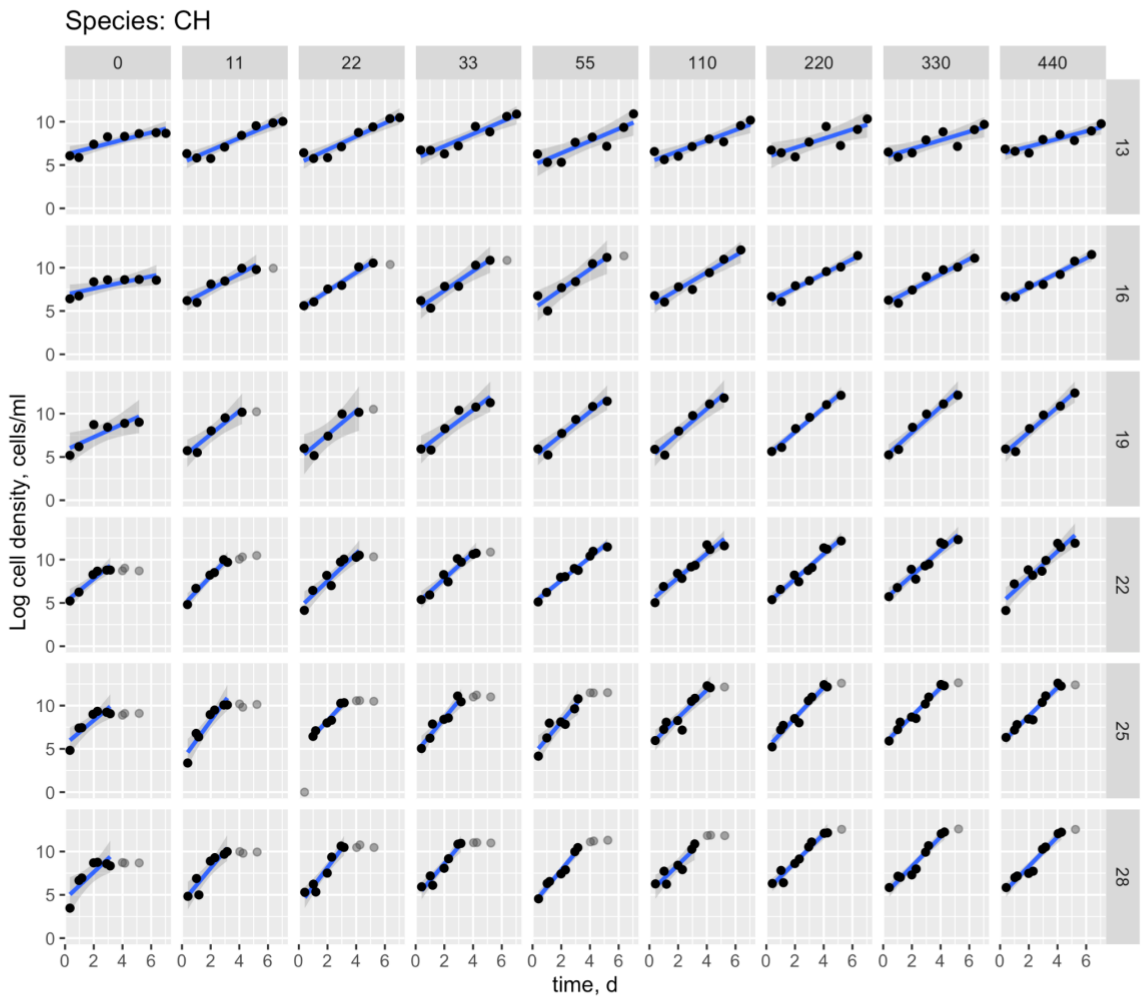


Fig. S5. Cell counts of *Chlamydomonas sp. (CH)* as a function of time. Grey data points indicate observations putatively outside the exponential growth phase, or deemed as outliers, and left out of linear model fits. Blue lines with shading show best-fit linear model with standard error to logged cell density, representing intrinsic rate of growth (*r*).

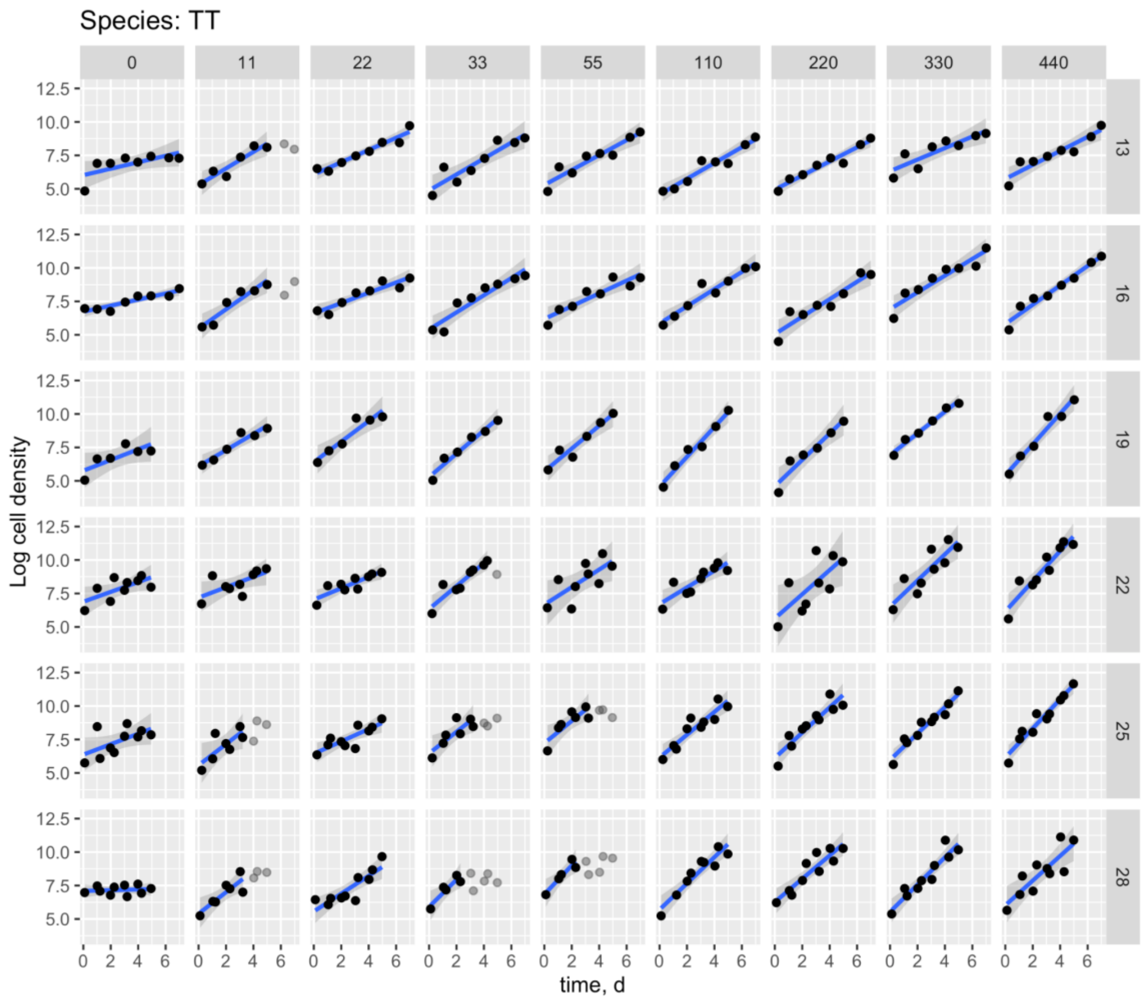


Fig. S6. Cell counts of *Tetraselmis tetrahele (TT)* as a function of time. Grey data points indicate observations putatively outside the exponential growth phase, or deemed as outliers, and left out of linear model fits. Blue lines with shading show best-fit linear model with standard error to logged cell density, representing intrinsic rate of growth (*r*).

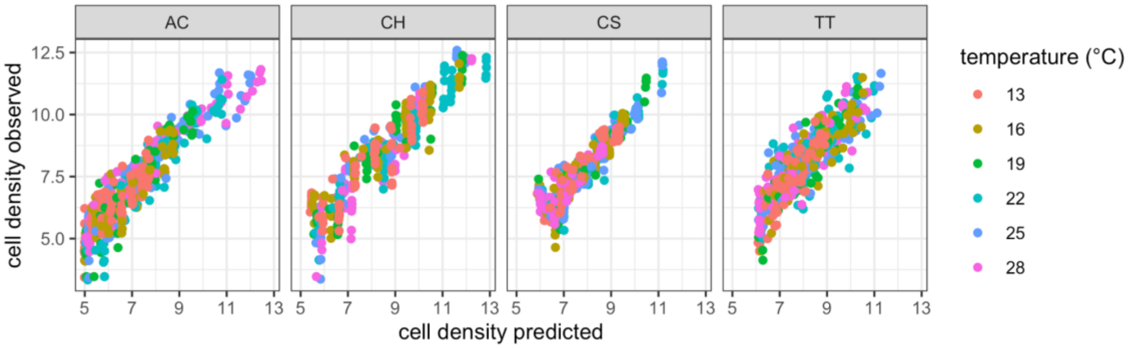


Fig. S7. Displays of fit for models of cell density as a function of time, temperature, and resource concentration in Experiment 1. Best-fit model predictions (x-axis) are from the Norberg Resource Model with varying Ks for species AC and TT, and from the Norberg Resource Model for species CH and CS. Species acronyms correspond to Table 2.

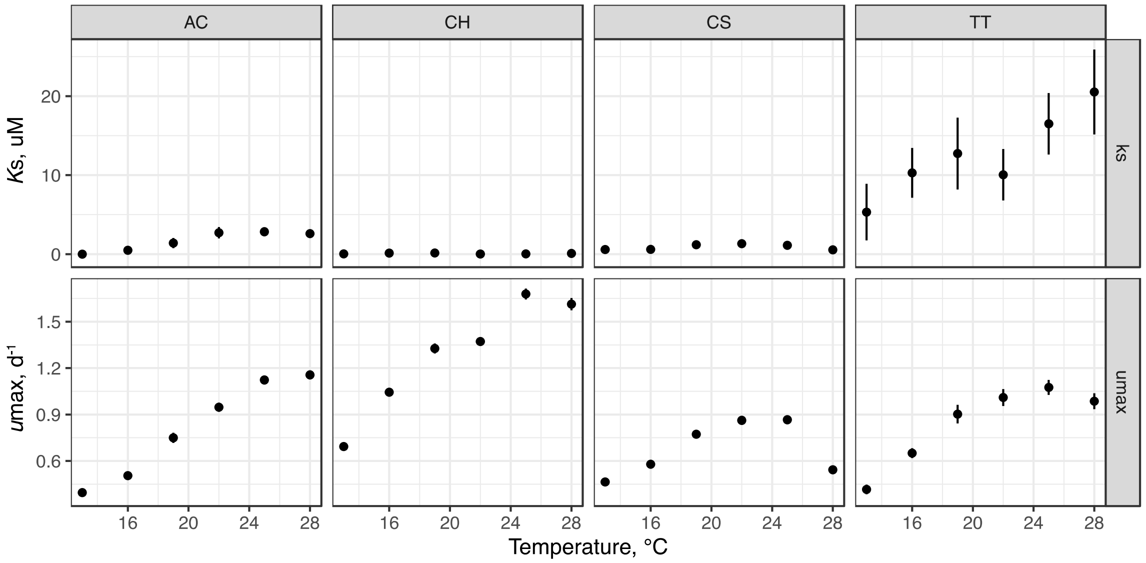


Fig. S8. Patterns of *Ks* and umax with temperature, from indirect model fitting within each temperature treatment. Species acronyms correspond to Table 2, errors represent ± SE. Growth rates at the highest nutrient concentration were used for umax.

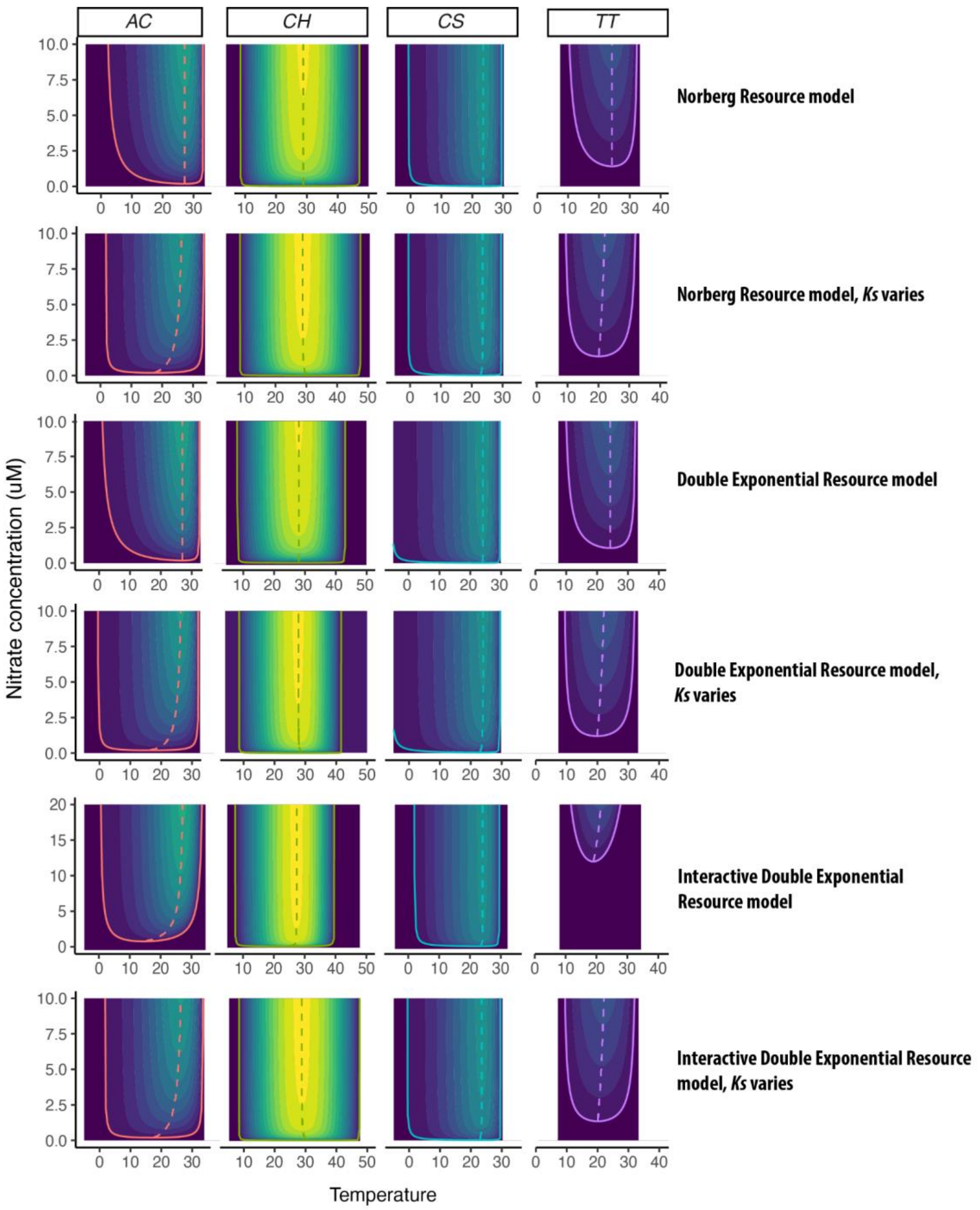


Fig. S9. Predicted growth rate, optimum temperature, and minimum resource requirement from each model. Color contours represent predicted growth rate as a function of temperature and nitrate concentration according to each model fit (rows, labeled on the right) for species (columns). Solid lines indicate minimum resource requirement ($R*$), and dotted lines indicate optimum temperature for growth (*Topt*). *Topt* is predicted to vary with temperature in models that allow Ks to vary with temperature, or in the interactive double exponential model with Ks fixed. Model names correspond to Table S2 and species acronyms correspond to Table 2.

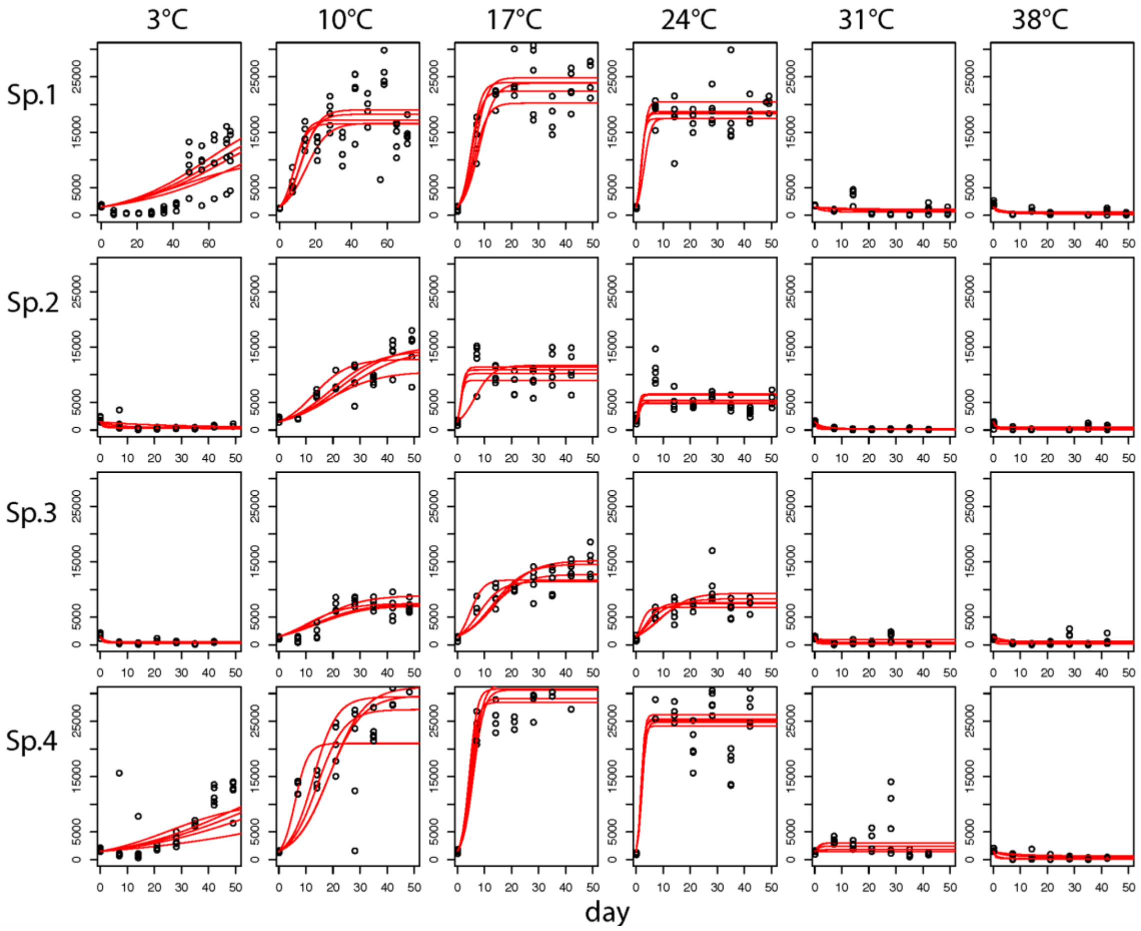


Fig. S10. Logistic growth models fit to cell density as a function of time in Experiment 2. For each species (rows) and temperature treatment (columns), points indicate cell density across 5 replicate cultures of each species maintained in semi-continuous flow, and red lines represent best-fit logistic growth model to each replicate.

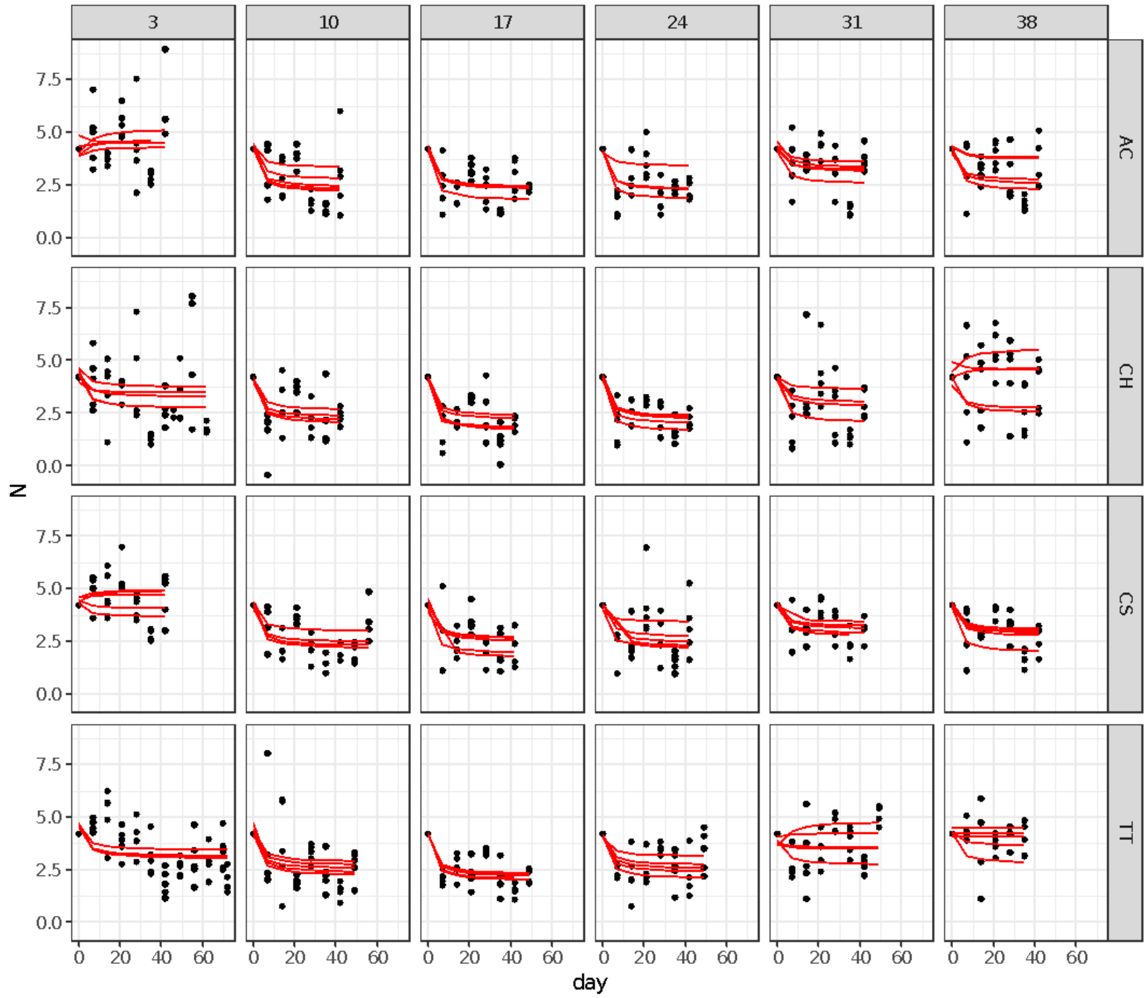


Fig. S11. Decay function fit to nitrate concentration as a function of time in Experiment 2. For each species (rows) and temperature treatment (columns), points indicate nitrate across 5 replicate cultures of each species maintained in semi-continuous flow, and red lines represent best-fit decay model fit to each replicate.

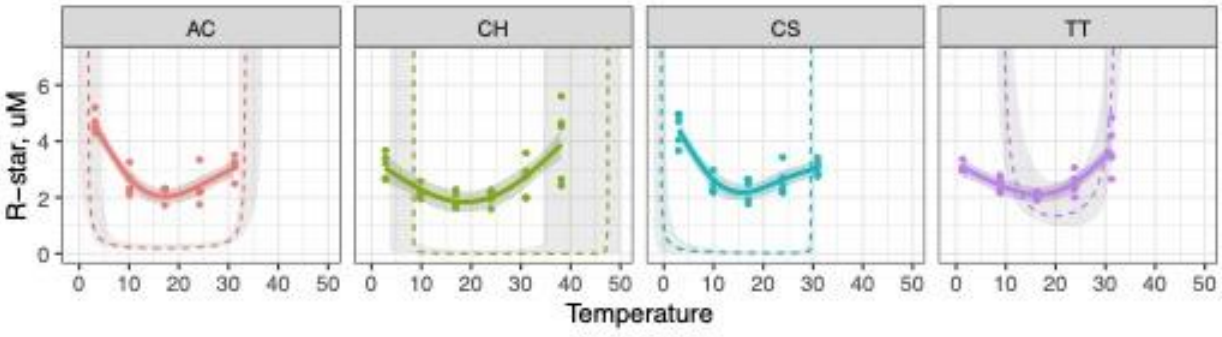


Fig. S12. Minimum resource requirements($R*$) from Experiments 1 and 2 displayed together. Dashed lines show the isocline of zero net-growth as a function of temperature from best-fit models to batch cultures in Experiment 1. Solid lines show GAM-model fits to equilibrium nitrate concentration in replicate continuous flow experiments across temperatures.

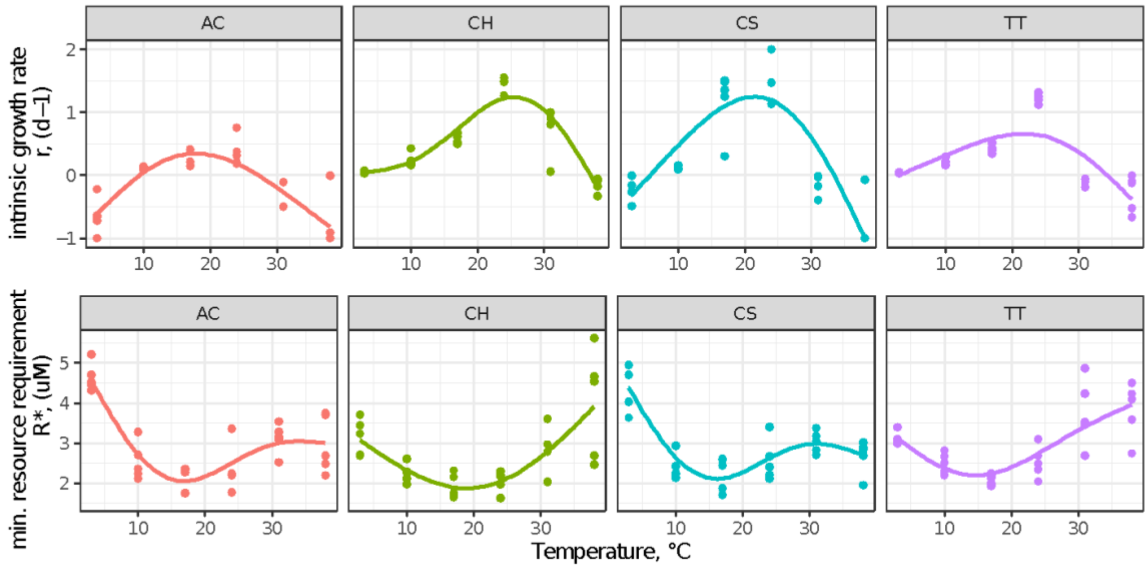


Fig. S13. Estimated growth rates (r) and minimum resource requirements (*R**) from Experiment 2 without exclusion of highest temperature. Population growth rates (r) were estimated from logistic-growth equations fit to cell density as a function of time; minimum resource requirements were estimated as asymptotic environmental nitrate concentrations at population equilibrium. Points indicate estimated values from each independent replicate, and lines with shading indicate best-fit generalized additive models.
